# CS - plasticity and behavioral-state integration in the posterior insula during auditory fear learning

**DOI:** 10.1101/2025.09.25.678609

**Authors:** Denys Osypenko, Shriya Palchaudhuri, Olexiy Kochubey, Ralf Schneggenburger

**Affiliations:** Laboratory for Synaptic Mechanisms, Brain Mind Institute, School of Life Science, Ecole Polytechnique Fédérale de Lausanne (EPFL), Switzerland; Department of Fundamental Neurosciences, University of Lausanne; Neuroscience Institute, New York University Grossman School of Medicine, New York, NY, USA; Centre for Brain and Mind, National Centre for Biological Sciences – TIFR, Bangalore, India

## Abstract

During fear learning, animals learn to associate sensory cues (conditioned stimulus, CS) with aversive outcomes, and neurons in several brain areas become entrained to the CS. The posterior insular cortex (pInsCx) contains representations of internal states and sensory cues, amongst them auditory stimuli. Nevertheless, the possibility of plastic encoding of auditory responses in the pInsCx during fear learning, and its underlying synaptic mechanisms have not been addressed. Using single-unit recordings in the pInsCx of male mice during fear learning, we find that ∼ 10% of putative principal neurons acquire a response to an auditory CS during fear learning (“CS learners”). CS learners are enriched in the insula auditory field (IAF) of the dorsal pInsCx, and strongly overlap with a larger neuronal subpopulation which increases its activity during movement initiation. Using optogenetic circuit mapping, we find that the IAF receives glutamatergic synapses from the non-lemniscal auditory thalamus and the auditory cortex (A1); fear learning induces a postsynaptic form of LTP at the cortical, but not at the thalamic input synapse. Combined single-unit recordings and optogenetic axon silencing showed that the acquired CS-response during fear recall depends on transmission at the A1 → pInsCx synapse. Our study shows that the pInsCx generates a plastic representation of an auditory CS during fear learning, driven by LTP of an associative cortical input. Future work should further investigate how the integration of CS- and movement information in the insula contributes to the expression of auditory-cued fear memories.

## Introduction

Aversively-motivated learning allows animals to learn about environmental cues that predict danger (Janak and Tye, 2015; Feinberg and Mallatt, 2017; Fanselow, 2018). Studying the neuronal circuits that enable aversively-motivated learning should allow us to learn about basic principles of brain function, as well as to obtain insight into the neural mechanisms that underlie neuropsychiatric disease like anxiety and post-traumatic stress disorders (Ressler et al., 2022; Ren et al., 2024). During fear learning, an auditory stimulus (conditioned stimulus, CS) acquires a negative emotional valence when paired with a mildly aversive outcome (unconditioned stimulus, US). Much work using these protocols has shown that the amygdala has major roles for the acquisition, consolidation and recall of an auditory-cued fear memory (for reviews, see LeDoux, 2000; Maren, 2001; Herry and Johansen, 2014; Tovote et al., 2015). Nevertheless, many other brain areas besides the amygdala are involved in fear learning (Dalmay et al., 2019; Szönyi et al., 2019; Barsy et al., 2020; Frontera et al., 2020; Taylor et al., 2021; Roy et al., 2022).

The insular cortex is conserved across mammalian species, and has been involved in various brain functions, ranging from the processing of sensory information to cognitive, emotional and autonomic functions (Nieuwenhuys, 2012; Gogolla, 2017). It is useful to sub-divide the insula into anterior, middle and posterior parts (Gogolla, 2017). An important function of the anterior-middle part of the insula is the processing of gustatory information and gustatory-related learning (Ogawa et al., 1990; Chen et al., 2011; Kusumoto-Yoshida et al., 2015; for a review see Maffei et al., 2012). More posteriorly, the insula processes somatosensory and nociceptive signals from the body surface both in humans and rodents (Rodgers et al., 2008; Coghill, 2020), and recent work in mice has revealed temperature coding in the posterior insular cortex (pInsCx)(Vestergaard et al., 2023). Furthermore, the pInsCx contains a representation of sensory signals arising from internal organs (Cechetto and Saper, 1987) and in turn can exert a top-down control over autonomic functions (Yasui et al., 1991); thus, the pInsCx is also referred to as visceral cortex (VISC; Allen Brain Atlas, 2014). Interestingly, a sub-region of the pInsCx in rats and mice was shown to process auditory information, and this area has been referred to as the “insular auditory field” (IAF) (Rodgers et al., 2008; Sawatari et al., 2011; Gogolla et al., 2014; Vestergaard et al., 2023). Together, these findings highlight multisensory integration in the insular cortex, as well as regional specificity of sensory modality processing mainly along the a-p axis of the insular cortex, but possibly also along dorso-ventral gradients.

Recent studies using optogenetic tools in the mouse model have started to investigate the function of various insular cortex subregions in anxiety-related behaviors, aversive state coding, and fear learning. One study showed that the pInsCx codes for aversive states but found no role for this area in fear learning (Gehrlach et al., 2019). Other studies have found that the anterior-middle region of the insula is involved in anxiety (Nicolas et al., 2023) and fear learning (Wang et al., 2022). We recently reported that the pInsCx contains a representation of aversive footshocks with both US-onset and US-offset responses, and that US-offset information is transmitted from the pInsCx to the lateral amygdala. Nevertheless, processing of footshock information by pInsCx principal neurons was not necessary for fear learning (Palchaudhuri et al., 2025). On the other hand, given that the pInsCx contains an auditory representation, it seems possible that sub-populations of neurons in the pInsCx acquire CS-responses during fear learning, and that this plasticity contributes to the formation, and/or retrieval of an auditory-cued fear memory (Rodgers et al., 2008).

Accordingly, we here use *in-vivo* single unit recordings in male mice, combined with *in-vivo*- and *ex-vivo* optogenetic techniques, to investigate whether the pInsCx elaborates a plastic CS representation during fear learning and if yes, to investigate its underlying synaptic mechanisms.

## Results

### An area at the pInsCx – S2 border shows plastic responses to tones during auditory-cued fear learning

To investigate how neurons in the pInsCx code for sensory stimuli and behavioral states during aversively-motivated learning, we performed single-unit recordings in the pInsCx of freely behaving mice during fear learning (Figure 1A-C). We employed a three-day fear learning protocol, with habituation- , training - and recall sessions (Figure 1A, Methods).

**Figure 1.**
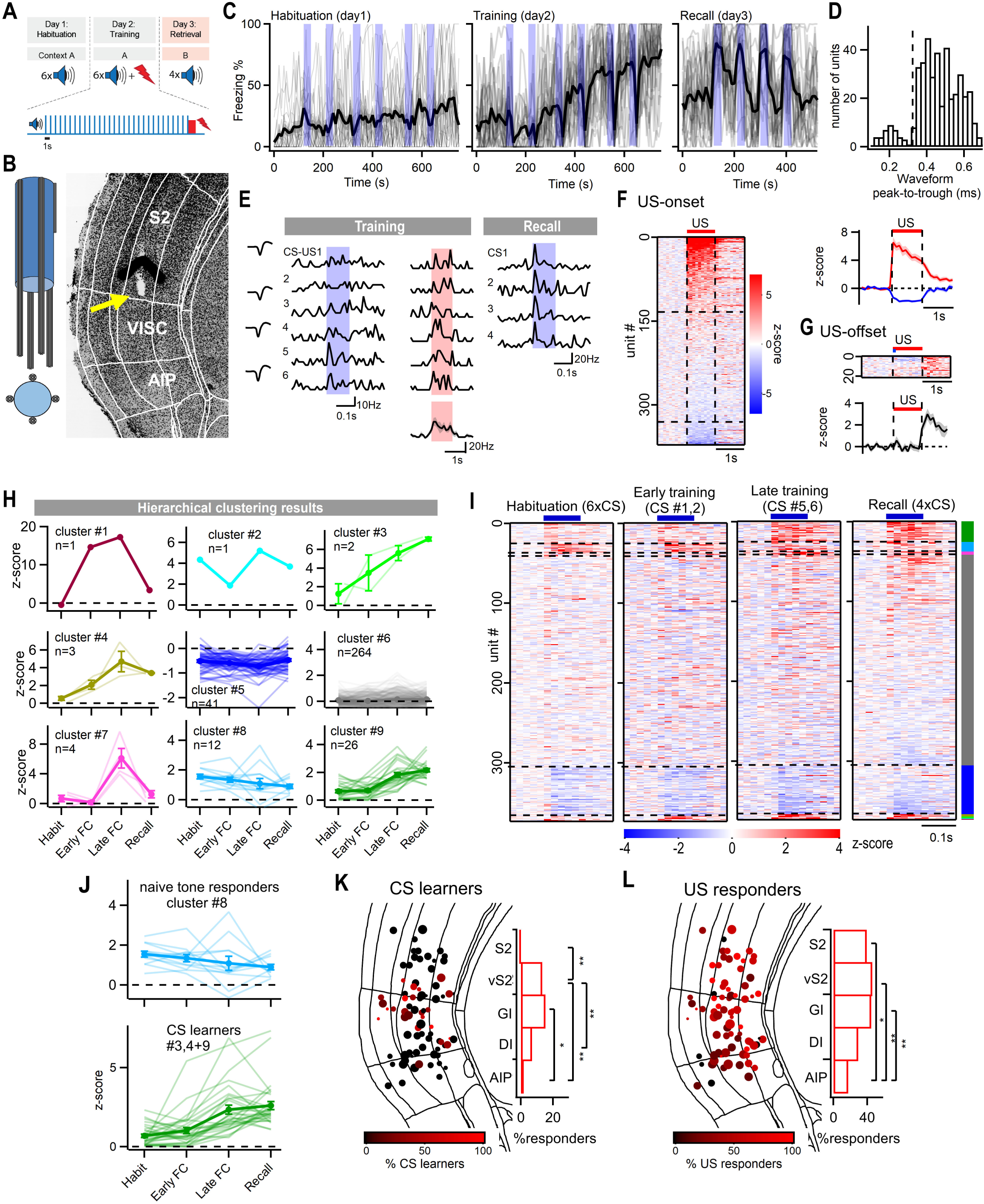
Single-unit CS and US responses in the pInsCx during a three-day fear learning protocol. ***A,*** Schematic of the auditory fear learning protocol. ***B,*** Schematic of an optrode assembly (*left*), and an example brain section for post-hoc localization of a tetrode recording tip (*right*; arrow). ***C,*** Individual (thin grey lines) and average (thick black lines) freezing curves (10 s time bin) of N = 20 mice from the *in-vivo* optrode recordings in Figures 1, 2. ***D,*** Histogram of the waveform width of all the recorded units (n = 420, in N=20 mice). Units with widths above- and below 0.35 ms were regarded as putative principal neurons (n = 374) and putative FSIs (n = 46), respectively. ***E,*** Example of a CS learner unit that also responded to the US. CS- (blue shading) and US-evoked (pink shading) AP activity during the training session (*left*) and the recall session (*right*). Insets on the left show the unit’s average waveform on each tetrode channel. The trace on the bottom shows the average US response. ***F,*** Raster plot showing the average PSTH for the US responses of all putative principal neurons (n = 374), sorted in decreasing order of z-score values. Dashed lines correspond to z-score values above and below which the units were classified as positive US-onset responders (*top*, n = 134) or US-inhibited units (*bottom*, n = 41), respectively. The inset on the right shows the average footshock-evoked activity of all positive US-onset responders (red trace) and of all US-inhibited units (blue trace). ***G***, Re-sorting the units for response maxima from 1-1.5 s after US-onset reveals n = 22 US-offset responders (*top*); the average trace is also shown (*bottom*). ***H,*** Results of unsupervised hierarchical clustering of CS-plasticity patterns observed across all putative principal neuron units (n=374). ***I,*** Raster plots of average CS-aligned activity of the units, sorted according to the cluster identity (see the color-coded bar, *rightt*). ***J,*** Individual units and average plasticity plots of cluster #8 (*top*, “Naive tone responders”) and of the pooled clusters #3, #4, #9 (bottom; “CS-learners”). ***K,*** *Left*, coronal view of the percentage of CS-learners on each tetrode, reconstructed from post-hoc histology data. *Right*, histogram showing the percentage of the CS-learners binned along five dorsoventral areas: S2, ventral S2 (vS2), pInsCS-GI (GI), pInsCx-DI (DI), pInsCx-AI (AIP). Note the predominant location of CS-learner units in the vS2 and GI. ***L,*** Data presentation as in K, now for the percentage of positive US-onset responders on each tetrode. For statistical parameters in K and L, see Results text.

During the training session, when each tone block (CS, conditioned stimulus) was followed by an aversive footshock, the mice displayed increasing levels of freezing. Furthermore, during fear memory recall, the mice increased the time spent freezing in response to each CS (Figure 1C; N = 20 mice). Thus, the mice displayed auditory-cued fear learning under conditions of *in-vivo* single-unit recordings.

Post-hoc histology and alignment to a mouse brain atlas showed that the recording electrodes were located in the granular- and dysgranular areas of the pInsCx, and in the dorsally adjacent S2 (Figure 1B; see also Figure 1K, L, below). We classified the recorded units as putative principal neurons, or as fast spiking interneurons based on their AP-waveform widths (n = 374 and 46 units respectively; N = 20 mice; Figure 1D). We will first present the responses of putative principal neurons to US- and CS stimulation during fear learning. We found that 36% of the units responded with an increased AP firing to footshocks presented as USs during the training session (Figure 1E, F). The US-response showed a tendency to increase in amplitude with repeated footshocks, with the first response being significantly smaller than the sixth (Figure S1). Another subpopulation of units responded with a *decrease* in AP-firing to the US (Figure 1F, *bottom*; 11 %). In addition, some units responded at the termination of the US (US-offset responses, Figure 1G; Palchaudhuri et al., 2025). Thus, a sub-population of principal neurons in the pInsCx processes aversive footshock information, as expected from previous studies (Rodgers et al., 2008; Gehrlach et al., 2019; Palchaudhuri et al., 2025).

We next investigated the responses of pInsCx neurons to the auditory stimuli (CS) presented throughout the fear learning protocol. To analyze the temporal dynamics unbiasedly, we separated the data into four time points (habituation session, early - and late training session, and recall session), and performed hierarchical clustering on session-averaged mean CS responses of individual units (Methods). At the depth of our clustering, there were nine different response types (Figure 1H). Interestingly, clusters 3, 4 and 9 showed an increasing CS-response throughout the fear learning protocol; we will refer to this response dynamics as “CS-learners” (Figure 1H-J; n = 31 / 374 units or 8.3%). Cluster 8 showed a nearly constant response to the tone stimuli over the course of fear learning; we refer to this response type as “Naive tone responders” (Figure 1H-J; n = 12 / 374 units or 3.1 %). Yet other sub-populations of pInsCx neurons showed constant negative responses to the CS, or else were non-responsive (Figure 1H, I; clusters 5 and 6). The overlap of CS- and US-responses is relevant for a mechanistic understanding of synaptic plasticity induced during fear learning (Herry and Johansen, 2014; Palchaudhuri et al., 2022). We found that amongst the n = 31 CS-learners, n = 19 showed an increased AP-firing in response to the US (see also Figure 2J), and none showed an inhibition of AP-firing induced by the US. Taken together, single-unit recordings in the pInsCx during a three-day fear learning protocol reveals neurons responsive to the footshock stimulation (US), as well as subpopulations of neurons that acquire a CS-response (CS-learners), and neurons responsive to tones throughout all sessions of the fear learning protocol (Naive tone responders).

**Figure 2.**
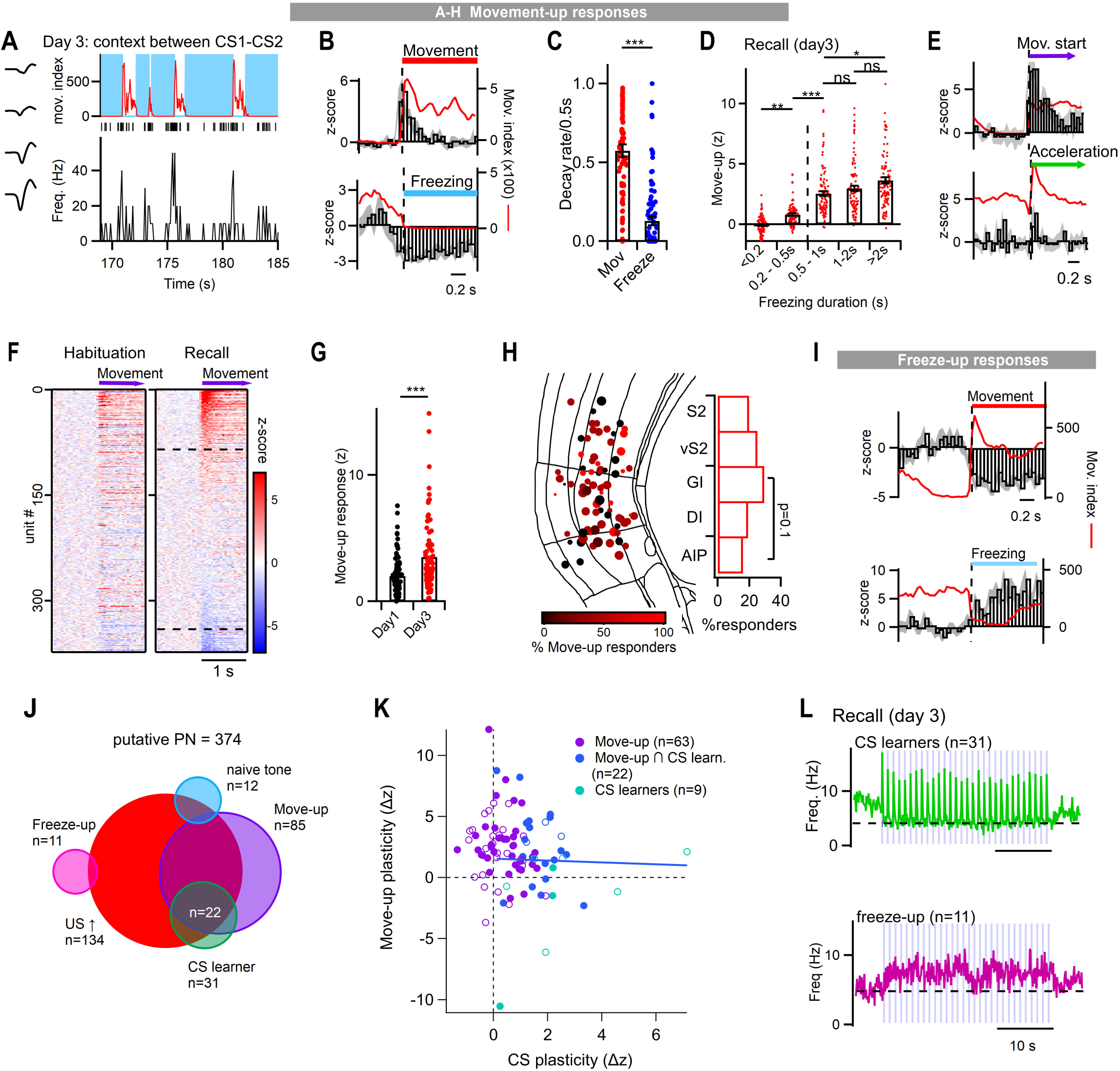
A robust movement-onset representation in the pInsCx. Data from n = 374 putative principal neurons from N = 20 mice are shown. ***A,*** Example of a unit active at the onset of movement, recorded in between CS1 and - 2 during the recall session. *Top*, movement trace (red trace, left axis) and freezing state of the mouse (blue). The average tetrode waveforms are shown on the left. *Bottom*, raster plot of AP-activity and AP-firing frequency of the unit. ***B,*** Black: average PSTH (50 ms bin width) of the activity of an example movement-up / freezing-down unit, aligned to the onset of movement (*top*) and to the onset of freezing (*bottom*). The red traces are the corresponding average movement traces of the mouse. ***C,*** Decay rates of the movement-up response and of the freezing-down responses (n = 85 units, p<0.001; Wilcoxon test). ***D,*** Amplitudes of the movement-up responses recorded during fear recall, sorted according to the duration of the preceding freezing episode. Note a consistently stronger movement-up response for the longer freezing episodes (see Results for statistical parameters). ***E,*** Another example of a movement-up responder, aligned to all movement-onset events (*top*), and to movement acceleration (*bottom*). Note the absence of consistent AP-firing in response to acceleration. ***F,*** Raster plots of the activity of all units aligned to movement-onset events during the habituation - (*left*) and recall session (*right*). The traces were sorted according to the response strength during the recall session; dashed lines show thresholds for movement-up (top line) and movement-down responses (lower line). ***G,*** Amplitudes of the average movement-up responses for each unit during the habituation session (*left*) and the recall session (right), analyzed for freezing episodes > 2 s (n = 85 units; p < 0.0001; paired t-test). ***H,*** Spatial distribution of the percentage of movement-up responders on each tetrode. ***I,*** Example unit of a freeze-up response aligned to the freezing to movement transitions (*top*), and to the movement to freezing transitions (*bottom*). ***J,*** Venn diagram summarizing the overlap of the various response types (n = 374 units from putative principal neurons; N = 20 mice). ***K,*** Scatter plot of the plasticity of the movement-up responses versus the CS-learner plasticity, for all movement-up and CS-learner units. The colors code for the overlap of the response types (*violet*; pure movement-up responders [n = 63]; blue: movement-up responders that were also CS-learners [n=22]; green: pure CS-learners [n=9]). Closed - and open data points identify units with, and without a positive US-onset response, respectively. The data was fitted by linear regression (blue line, r = - 0.028, p = 0.88; Spearman correlation test). ***L,*** Average CS-evoked AP-firing responses during the fear recall session of the CS-learners (*top*, green trace, n = 31 units), and of the freeze-up responders (*bottom*, n = 11 units). See also Figure S2 for part of the corresponding data for FSIs.

Using post-hoc histological analysis of the tetrode positions, we next correlated the position of the recording tetrodes with the fraction of CS-learners and US responders. This showed that tetrodes implanted in the granular part of the pInsCx (pInsCx-GI), and in the ventral S2 (vS2) sampled a significantly higher percentage of CS learner units than tetrodes at other locations (Figure 1K; vS2 vs S2, pInsCx-DI and AIP; p < 0.01; pInsCx-GI vs AIP, p < 0.05; Fisher’s exact test with Bonferroni correction). Regarding the US responses, electrodes in the agranular part of the pInsCx (AIP) contained a significantly smaller fraction of US-responses than electrodes in the other areas (Figure 1L; AIP vs pInsCx-GI and vS2, p < 0.01; AIP vs S2, p < 0.05; Fisher’s exact test with Bonferroni correction). Taken together, these findings show that a sub-population of neurons in the pInsCx acquires a response to an aversively-motivated auditory CS during fear learning. The CS-learner units were enriched in the pInsCx-GI and in the vS2, and we assume that this region corresponds to the previously described IAF (Rodgers et al., 2008; Sawatari et al., 2011; Gogolla et al., 2014).

### A subpopulation of pInsCx neurons codes for movement-onset

During fear learning, mice show increasing levels of freezing, and freezing is occasionally interrupted by movement re-initiation (Figure 1A; Figure 2A). To test whether the activity of pInsCx neurons correlates with transitions between freezing and movement, we aligned the activity of the recorded units to both types of transitions detected in each mouse. This revealed a sub-population of neurons with increased AP-firing upon movement re-initiation after freezing (Figure 2A, Figure 2B, *top*; n = 85 / 374 or 23%). In turn, during freezing, these neurons showed a *decrease* of their AP-firing rate (Figure 2A; Figure 2B, *bottom*), thus characterizing this response as a movement-up / freezing-down response. The movement-up response was transient, and had a significantly faster decay rate than the freezing-down response (Figure 2C, p < 0.001, Wilcoxon test). Interestingly, the amplitude of the movement-up responses depended on the duration of the preceding freezing bout, with maximal responses attained for freezing bouts longer than ∼ 1 s (Figure 2D; n = 85 units, p < 0.001, repeated measures ANOVA followed by Tukey’s post-hoc tests; <0.2s vs 0.2 - 0.5 s, p < 0.01; 0.2 - 0.5s vs 0.5 - 1s, p < 0.001, and 0.5 - 1s vs >2s, p < 0.05; all other comparisons not significant). Furthermore, these units showed only a small response to acceleration, suggesting that they primarily code for behavioral transitions rather than for movement velocity (Figure 2E).

To assess whether the movement-up responses were stable, or else were modulated in a plastic manner during the course of fear learning, we compared their amplitudes between the habituation - and the recall sessions for all units. This revealed that the movement-up responses had increased amplitudes during the recall session as compared to the habituation session (Figure 2F, G; p <0.001, paired t-test). Neurons with movement-up / freezing-down responses did not show a significant spatial clustering within subdomains of the S2 and pInsCx (Figure 2H). In addition to movement-up responses, we found a smaller percentage of pInsCx neurons with the opposite movement responses, i.e. their AP-firing *decreased* during movement, and increased during freezing (“freeze-up” responses; Figure 2I; 29/374 or 7.8% of all units; see also Gehrlach et al., 2019). The *in-vivo* single-unit recordings thus reveal that beyond responses to the CS and US, there are neurons that code for the movement state of the animal, with a relatively large sub-population of movement-up responders (23%), and a smaller sub-population with freeze-up responses (Figure 2J). Importantly, a large fraction of the CS learners (22 out of 31) showed movement-up / freeze-down response (Figure 2J).

We found above that the movement-up responses were increased after fear learning, and that a majority of the CS learner units also showed movement-up responses. We therefore next analyzed whether the degree of plasticity of the CS-representation, and of the movement-up representation was correlated. For this, we plotted the amplitude of the plasticity of movement-up responses, as a function of the CS-plasticity, for the sum of movement-up responders and CS-learners (n = 94 units). In the resulting plot, most data points were localized in the top-right quadrant, which suggests overall positive plasticity of both the CS-representation, and the movement-up representation, in-keeping with the above results. A line fit restricted to the CS learners returned a slope close to zero (Figure 2K, blue and green data points; r = 0.028, p = 0.88, Spearman correlation test); a line fit to all data points returned a negative correlation (r = -0.227, p = 0.028; not shown). We conclude that the CS-plasticity, and the plasticity of the movement-up representation was not strongly correlated.

Because many CS-learners also show a movement-up / freezing-down response, one might expect that during fear recall, when mice show a high amount of freezing, the AP-firing response of the CS-learner neurons is shaped by two factors: First, by transient responses to each tone beep, and second, by a tonic *decrease* in AP-firing caused by the effect of freezing on the activity of these neurons. Indeed, averaging the AP-firing responses of all CS-learners during the recall session revealed phasic, short-latency responses to individual tone beeps, followed by a tonic decrease in AP-firing (Figure 2L, *top*; n = 31 units). On the other hand, as expected, units with a freeze-up response showed a tonic AP-firing increase during the CS at fear memory recall, but phasic tone-driven responses were not visible (Figure 2L, *bottom*).

Thus, the AP-firing response to the CS of movement-state responsive neurons is influenced by the behavior of the mice especially during the recall session, when the mice display learned freezing in response to the CS. Thus, an overlap of sensory-driven responses with movement-state responses can result in apparent sensory-driven activity, which should be considered when analyzing in-vivo responses in freely behaving animals.

We also analyzed the data for n = 46 recorded putative FSIs, based on their short AP-waveform (< 0.35 ms; see Figure 1D). We observed nearly all response types that we also found in putative pyramidal neurons, with an over-representation of naive tone responders in the FSI as compared to the pyramidal neurons (Figure S2). Taken together, *in-vivo* single unit recordings identify putative principal neurons with different response - and plasticity types in response to the two sensory stimuli presented during fear learning (CS and US), and with a strong coding for movement onset after aversively-motivated freezing. Furthermore, during fear memory recall, the responses of CS learner units is determined by a mix of sensory - and behavioral state information.

### Auditory thalamus sends a robust glutamatergic projection to the pInsCx

We next investigated the synaptic mechanisms that underlie the plasticity of CS-responses in the pInsCx. We hypothesized that the plastic representation of a CS in the pInsCx is driven by long-term potentiation (LTP) at excitatory synapses that carry auditory information to the pInsCx. A previous study has investigated the connectivity of the IAF with the medial geniculate body (Takemoto et al., 2014). We wished to obtain independent evidence for brain areas that can carry auditory information to the pInsCx, and therefore performed retrograde labelling experiments (Figure 3A). To allow an unambiguous identification of the pInsCx relative to the S2, we employed Scnn1a^Cre^ x tdTomato mice; *Scnn1a* is a marker gene for layer 4 neurons in sensory cortices (Madisen et al., 2010). We found that tdTomato expression was abundant in layer 4 of the S2, and then terminated at the ventral end of S2, thus providing a molecular ruler for the pInsCx - S2 border (Figure 3B, arrow; Bokiniec et al., 2022; Palchaudhuri et al., 2025). We injected retrograde label (CTB) conjugated to either Alexa-488 or Alexa-647 at two dorsal-ventral levels, and observed back-labelled neurons in various forebrain areas (Figure 3). The two injections revealed separate, topologically arranged pools of back-labelled neurons in the contralateral pInsCx (Figure 3B, *right*).

**Figure 3.**
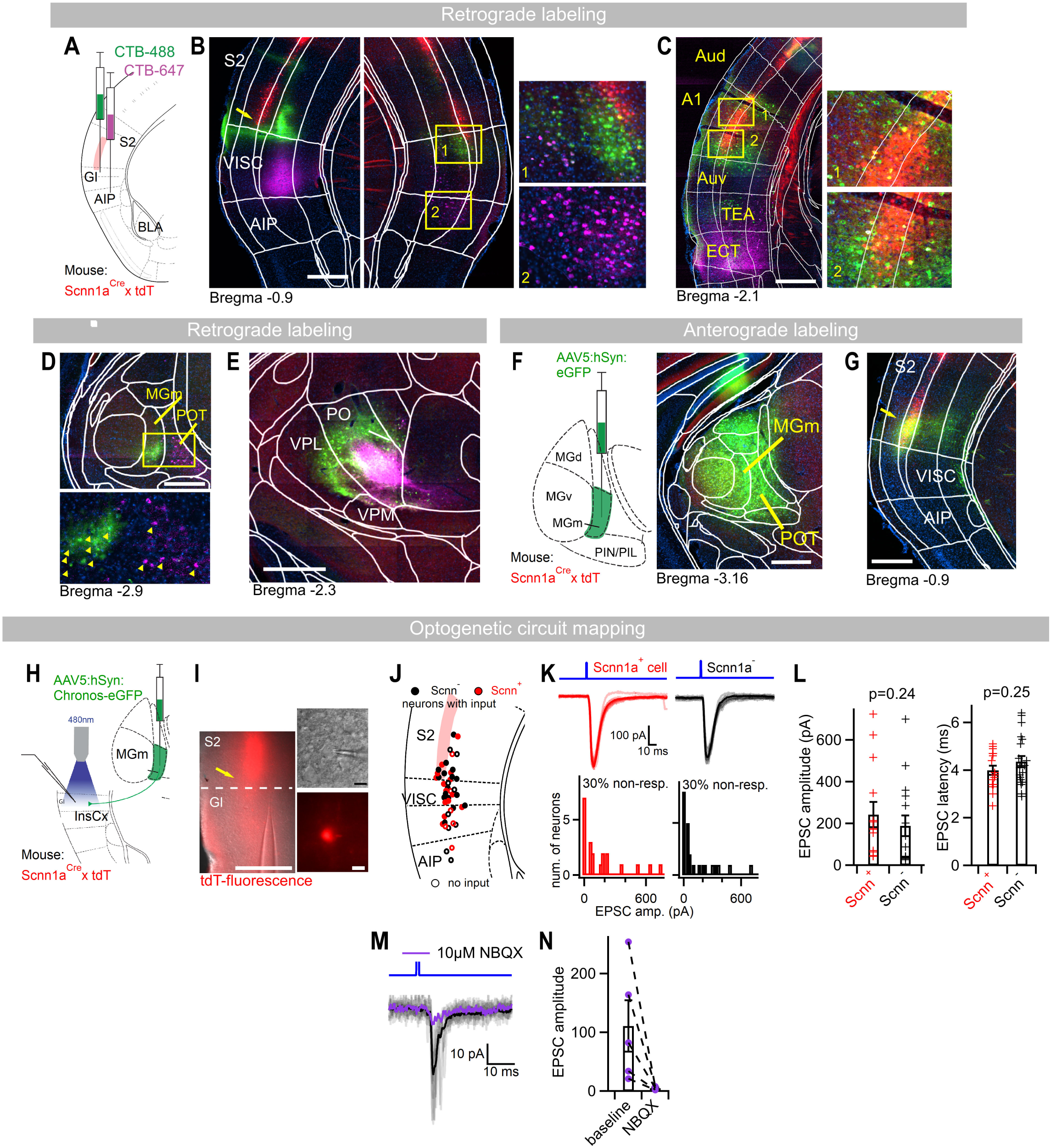
The pInsCx receives a robust glutamatergic synaptic input from the MGm. ***A,*** Schematic of the double-retrograde labeling experiment employing *Scnn1a^Cre^* x *lsl-tdTomato* mice to label layer 4 neurons, and injections of CTB-488 and CTB-647 into the pInsCx - S2 border, and into the AIP, respectively. ***B,*** Images of the CTB injection sites (*left*) and of retrogradely labelled neurons in the contralateral pInsCx (*right*). The insets on the *right* are higher-magnification images corresponding to the yellow boxes. Here, and in panels C - E, the green -, magenta- and red channels correspond to CTB-488, CTB-647and tdTomato. ***C,*** Example image of back-labelled neurons in the ipsilateral primary auditory cortex (A1) and more ventral cortical areas. The insets (*right*) show higher magnification images at the position of the two yellow boxes. Note the predominant back-labelling of neurons in A1 from the pInsCx - S2 border (CTB-488, green channel), and of neurons in the ECT from the AIP (CTB-647, magenta channel). ***D,*** Image of retrogradely labelled neurons in the auditory thalamus (*top*), and magnified image corresponding to the yellow box in the top panel (*bottom*; yellow arrowheads identify single back-labelled neurons).***E,*** Back-labelled neurons at the level of the somatosensory thalamus. ***F,*** *left*, schematic of the anterograde labeling experiment from the MGm. *Right*, image of the injection site showing eGFP fluorescence (green channel). ***G***, Example image showing eGFP-positive axons from the MGm (green channel). Note the tdTomato+ (Scnn1aCre+) band of layer 4 neurons, which identifies the pInsCx - S2 border (red channel; see arrow). ***H,*** Schematic of optogenetic circuit mapping experiment performed in *Scnn1aCre* x *tdT* mice. ***I***, Fluorescent image of an acute coronal brain slice during the patch-clamp experiment (*left*; red, tdTomato), and higher-resolution images of a tdTomato+ neuron being recorded (*right*). ***J,*** Map of recorded neurons that showed-, or did not show optogenetically-evoked EPSCs (filled, or open symbols, respectively). Red- and black symbols identify Scnn1a^Cre^-positive (i.e. tdTomato-+) or - negative neurons, respectively. ***K,*** *Top*: example recordings (overlay of 10 consecutive sweeps) of optically-evoked EPSC from a Scnn1a^Cre^-positive -, and negative neuron (red and black traces). *Bottom*, histograms of EPSC peak amplitudes measured in Scnn1a-positive and negative neurons (n = 14 and 16, respectively, from N = 4 mice). The first bin at zero sums up all the non-responsive cells. ***L,*** Amplitudes- and latencies of the EPSCs showed no difference between Scnn1a-positive and negative neurons (p = 0.24 and 0.25, t-test). ***M,*** Average- and individual EPSC traces before (grey and black) and after (violet) bath application of 10 μM NBQX in an example recording. ***N,*** NBQX strongly reduced the amplitude of the EPSCs (n=5 recordings in one mouse). Scale bars are 500 μm in B-G; I, *left*, and 10 μm in I, *right*.

Furthermore, neurons back-labelled from the pInsCx - S2 border (CTB-488; green channel) were found in the A1 (Figure 3C), in the medial part of the medial geniculate nucleus (MGm; a non-lemniscal part of the auditory thalamus; Figure 3D), as well as in the ventral posterolateral nucleus of the thalamus (VPL) and the Posterior complex of the thalamus (PO; Figure 3E), and in further brain areas (not shown). On the other hand, neurons back-labelled from the more ventral pInsCx-DI (CTB-647; magenta channel) were found in the Ectorhinal area (ECT), in the posterior triangular thalamic nucleus (POT), and in the PO (Figure 3C-E, magenta channel). These results from retrograde labelling identify two brain areas that might carry auditory information to the pInsCx: First, the non-lemniscal auditory thalamic nucleus MGm (see also Takemoto et al., 2014), and second, the primary auditory cortex, A1.

We next tested whether the MGm provides a functional, presumably glutamatergic connection to the pInsCx, using anterograde labelling and optogenetic circuit mapping (Petreanu et al., 2007; Little and Carter, 2012). Intriguingly, anterograde labelling with injections of AAV5:hSyn:eGFP into the auditory thalamus of Scnn1a^Cre^ x tdTomato mice (Figure 3F) produced a band of eGFP-expressing axons at the pInsCx - S2 border (Figure 3G, *arrow*); the location of the axonal bundle coincides with the location of CS-learners in the *in-vivo* recordings (Figure 1K). Next, employing optogenetic circuit mapping with expression of Chronos in the MGm (AAV5:hSyn:Chronos-eGFP), followed by recordings of neurons in the pInsCx, we observed robust blue-light evoked EPSCs (Figure 3H-L). We recorded neurons in layers 4 and 5 of the pInsCx (both Scnn1a+ and Scnn1a-neurons), sampling a dorsal-ventral spread of neurons in the pInsCx and vS2 (Figure 3J). We found that the EPSC amplitudes were indistinguishable between Scnn1a+ and - neurons (Figure 3J-L; p = 0.23; Mann-Whitney U-test). Interestingly, there was a higher density of connected neurons in the pInsCx-GI and vS2, as compared to more ventral recordings sites (Figure 3J), a finding which agrees with the location of afferent axons in the anatomical data (Figure 3G1). The optogenetically-evoked EPSCs had short latencies of ∼ 4 ms, and were blocked by the AMPA-receptor antagonist NBQX (10 µM; Figure 3M, N). These data show that the MGm makes a glutamatergic synapse onto neurons of the pInsCx-GI, and the adjacent ventral S2.

### A1 also provides a glutamatergic projection to the pInsCx

We found above that besides the MGm, the primary auditory cortex (A1) was also labelled in retrograde tracer experiments from the pInsCx (Figure 3C). Therefore, we next employed anterograde labelling and optogenetic circuit mapping to investigate whether the A1 similarly makes a glutamatergic input synapse to the pInsCx. Anterograde labeling with an eGFP - expressing viral vector injected into the A1 (AAV8:hSyn:eGFP) revealed a band of eGFP - positive axons at the pInsCx - S2 border (Figure 4A, B). Simultaneous anterograde labelling with a two-color approach (see Methods) showed that the inputs from the MGm and A1 are both localized in an overlapping area at the pInsCx - S2 border. Nevertheless, the two inputs target different cortical layers: The MGm input mainly terminated in layer 4, whereas the A1 input terminated in layer 2/3 and - 5 of the insular cortex (Figure 4C, D).

**Figure 4.**
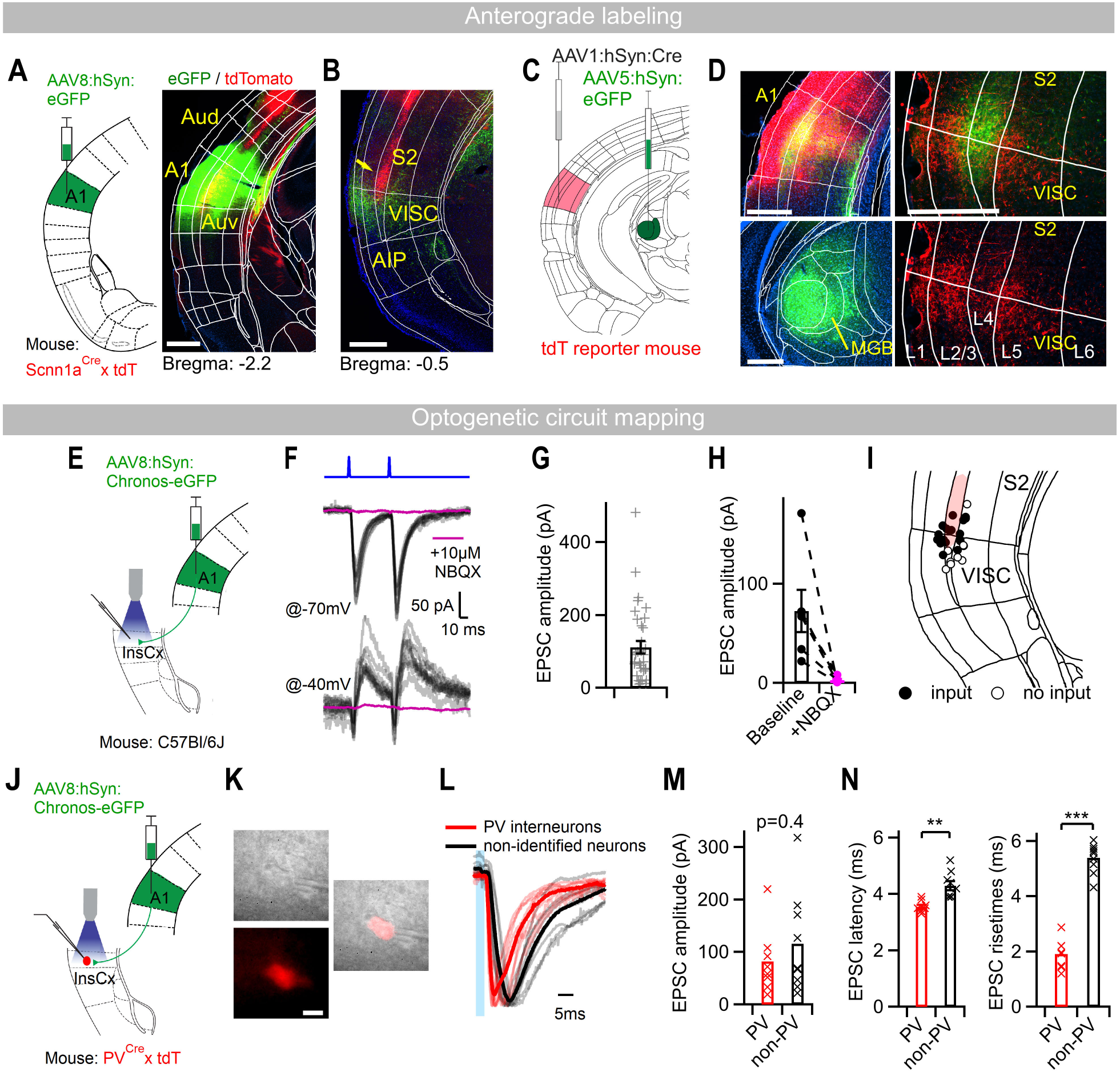
The pInsCx also receives a glutamatergic synaptic input from A1. ***A,*** *Left,* schematic of the anterograde labeling experiment from A1. *Right*, image of the injection site in the A1 (green channel: eGFP; red channel: tdTomato, i.e. Scnn1aCre+ neurons). ***B***, Image of the pInsCx showing eGFP+ axons from A1 (green channel) close to the pInsCx - S2 border, as indicated by the Scnn1a+ neurons (tdTomato; red channel; arrow). ***C,*** Schematic of dual anterograde labeling of A1 - and MGm projections, using a Cre-expressing vector in tdTomato reporter mice, and an eGFP-expressing vector in the MGm. ***D,*** *Left*: images of the injection sites in A1 (*top*) and MGm (*bottom*). *Right*: images of the pInsCx - S2 border. Note the tdTomato-positive axons that originate from the A1 in layer 2/3 and -5 (red channel), and the eGFP+ axons originating from the MGm in layer 4. ***E,*** Schematic of an optogenetic circuit mapping experiment at the presumed A1 → pInsCx connection. ***F,*** optogenetically-evoked EPSCs in a pInsCx neuron recorded at -70 mV (*top*) or, in another recording, at -40 mV. NBQX (10 µM) was applied in both recordings. For control conditions, ten subsequent EPSCs are shown (grey traces), whereas in the presence of NBQX an average of n = 10 traces is shown (violet trace). ***G,*** Individual and average EPSCs amplitudes recorded at -70 mV (n = 35 recordings from N = 4 mice). Cells not receiving a detectable EPSC are not included in the plot. ***H,*** Individual and average EPSCs peak amplitudes before (*left*) and after application of 10 µM NBQX (right; n = 6 recordings from N = 2 mice). ***I,*** Locations of the recorded neurons (see E-H) with - or without detectable input from the A1 (filled - and open symbols, respectively). ***J,*** Schematic of the optogenetic circuit mapping experiment employing *PV^Cre^ x tdTomato* mice to identify PV-interneurons in slice recordings. ***K,*** Transmission - and fluorescence images (red, tdTomato) of a recorded PV-interneuron. ***L,*** Peak-normalized EPSC traces recorded in tdTomato+ PV-interneurons (red; n=8) and in tdTomato- recorded neurons (grey; n=9). Thin and thick lines are individual - and average traces, respectively (n = 8 and 9). ***M,*** Average EPSC amplitudes recorded in tdTomato+ PV-interneurons (*left*, red symbols) and in tdTomato- neurons (*right*; n = 8 and 9 recordings). ***N,*** EPSC latencies (*left*) and EPSC risetimes (*right*) for the same recordings as shown in panels L and M. For statistical parameters, see Results. Scale bars are 500 μm in A, B and D, and 10 μm in K.

In optogenetic circuit mapping following the expression of Chronos in A1 (Figure 4E), we observed robust EPSCs elicited by blue-light pulses, which were blocked by NBQX (10 µM; Figure 4E-H; 95 ± 2% block, n = 6 cells). In a subset of recorded neurons we investigated whether the A1 input recruits a feedforward inhibition, by recording postsynaptic currents at - 40mV with a low [Cl^−^] pipette solution (8 mM; see Methods). A majority of the neurons recorded for this purpose (n = 7/8) showed a sequence of inward EPSCs, followed by delayed outward postsynaptic currents (Figure 4F, *bottom*). Both the inward- and the outward currents were blocked by NBQX (10 µM), suggesting that excitatory afferents from A1, besides connecting directly to principal neurons, engage a disynaptic feedforward inhibition of the pInsCx. To investigate the interneuron type involved in this feedforward inhibition, we performed additional experiments with PV^Cre^ x tdT mice, to target the recordings to PV-interneurons based on their tdTomato - fluorescence (Figure 4J, K). Consistent with the notion of feedforward inhibition, we observed short-latency optogenetically-evoked EPSC in PV-interneurons (Figure 4L-M). The amplitude of the EPSCs was not significantly different between PV-interneurons and non-labelled putative principal neurons recorded in the same slices (Figure 4M, p = 0.39; t-test). The latency and rise-time of the EPSCs were significantly shorter in PV-interneurons than in putative principal neurons (Figure 4N; p < 0.01 and p < 0.0001, t-test), as expected (Hu et al., 2014). Taken together, anterograde tracing and optogenetic circuit mapping reveal a functional synaptic input from A1 to the pInsCx, which additionally engages a feedforward inhibition via PV-interneurons.

### Fear learning drives LTP at cortical but not thalamic afferents to the pInsCx

We found that a subpopulation of neurons at the pInsCx - S2 border acquires auditory CS responses during fear learning, and that neurons in this area receive glutamatergic inputs from two auditory areas, the MGm and the A1 (Figures 1 - 4). We hypothesized that fear learning might drive long-term potentiation (LTP) at one, or both of these input synapses to the pInsCx, and that this plasticity might underlie the acquisition of CS-responsiveness during fear learning, that we observed in a sub-population of pInsCx neurons (CS-learners, Figure 1). Previous work has shown that measurements of the AMPA/NMDA ratio, and/or paired-pulse ratio (PPR) at a given synaptic connection after a behavioral learning experience, can assess postsynaptic and/or presynaptic forms of long-term plasticity (McKernan and Shinnick-Gallagher, 1997; Lucas et al., 2016; Kim and Cho, 2017; Rich et al., 2019; see Palchaudhuri et al., 2022 for a review). Therefore, we next measured the AMPA/NMDA ratio, and the PPR at the cortical- and thalamic inputs to the pInsCx after fear learning, to assess the possibility that fear learning causes pre- or postsynaptic forms of LTP at these synapses.

We first performed AMPA/NMDA ratio measurements at the MGm to pInsCx synapse. Mice were injected bilaterally into the MGm with an AAV vector driving the expression of Chronos (AAV5:hSyn:Chronos-eGFP; Figure 5A, B). Three to five weeks later, mice were randomly assigned to one of two groups: One group underwent standard fear learning (“CS+US” group), and a control group underwent a protocol without footshocks during the training session (“CS only” group; Figure 5A). Mice in the CS+US group displayed fear behavior characterized by an increasing freezing response during the training session, and by CS-driven freezing during recall on the background of a moderate level of context freezing (Figure 5C, red average traces, N = 7 mice; see also Mombelli et al., 2024; Palchaudhuri et al., 2025). Conversely, mice in the CS-only group showed minimal freezing throughout all sessions, as expected (Figure 5C, black traces; N = 6 mice).

**Figure 5.**
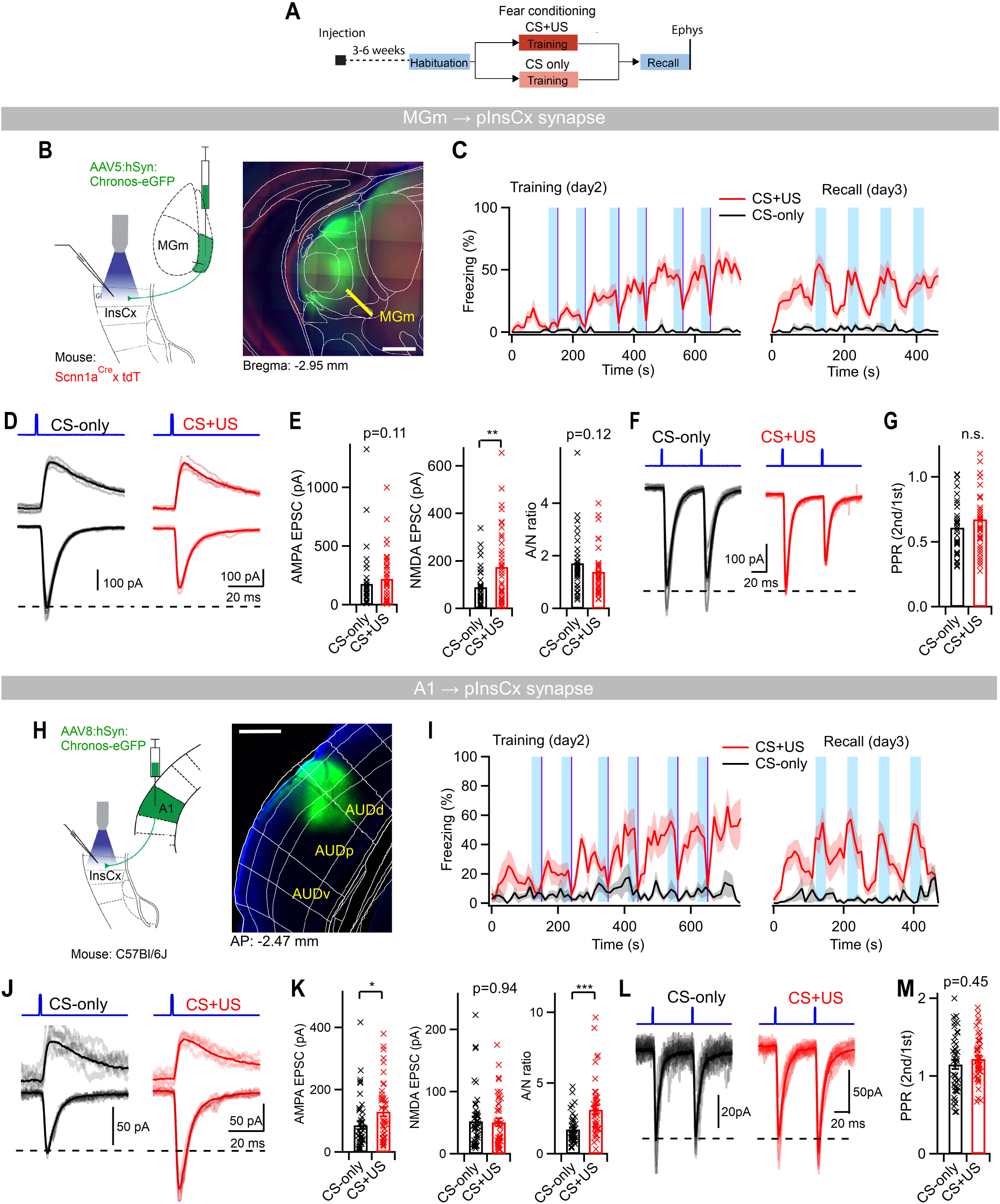
Fear learning drives canonical LTP at the cortical but not the thalamic afferents to the pInsCx. ***A,*** Timeline of the experimental approach for the experiments in this Figure. ***B***, *Left*, schematic of expression of Chronos in the MGm → pInsCx pathway. *Right*, Image of the auditory thalamus showing the expression of Chronos-eGFP at the injection site (green channel). ***C***, average freezing traces (shading is ±s.e.m.) in mice from the “CS+US” group (red; N = 7 mice) and from the CS-only group (black; N = 6) during the training - and recall sessions as indicated. ***D,*** Example single (grey or pink) and average EPSCs (black or red) measured at -70 mV (*bottom*), and at +50 mV (*top*), obtained from a CS-only mouse and a CS+US mouse (*left* and *right* panels, respectively). ***E,*** Individual and average values of the AMPA-EPSC amplitude (*left*), of the NMDA-EPSC amplitude (*middle*), and of the AMPA/NMDA ratio (*right*), measured in the two experimental groups at the MGm → pInsCx synapse. For statistical parameters, see Results. ***F,*** Example EPSCs recorded with a PPR protocol (interval, 50 ms) for the CS-only group (*left*) and the CS+US group (*right*). ***G,*** Individual and average values of PPR in both groups. ***H,*** Scheme showing the approach for expressing Chronos at the A1 → pInsCx pathway (*left*), and image of the auditory cortex showing the expression of Chronos-eGFP at the injection site (green channel, *right*). ***I,*** Similar data as shown in panel C, but now for the mice in the CS+US and CS-only group that expressed Chronos at the A1 → pInsCx pathway. ***J,*** Example EPSCs recorded at the A1 → pInsCx connection. The display is analogous to the one in panel D. ***K,*** Average and individual values of AMPA-EPSCs (*left*), NMDA-EPSCs (*middle*) and of the AMPA/NMDA ratio (*right*) quantified for the two behavioral groups, here measured at the A1 → pInsCx connection. ***L,*** Example EPSCs recorded with a PPR protocol for both behavioral groups, measured at the A1 → pInsCx synapse. ***M,*** Individual and average values of PPR in both behavioral groups. Scale bar, 500 µm in B and H.

Following the fear memory recall test, which was performed to assess whether individual mice showed fear memory expression, optogenetically-evoked EPSCs at the MGm to pInsCx connection were measured. The absolute AMPA-EPSCs amplitude (measured at -70 mV), and the AMPA/NMDA ratio were unchanged between the CS-only and CS+US groups (Figure 5D, E; p = 0.11 and p = 0.12, respectively, Mann-Whitney test; n = 39 cells recorded in N = 7 mice and n = 36 cells recorded in N = 6 mice for the CS+US - and CS-only groups, respectively). Interestingly, the NMDA-EPSCs (measured at + 50 mV) were larger in the CS+US group than in the CS-only group (p = 0.0036; Figure 5E), which might indicate a form of long-term plasticity of the NMDA-component at the MGm to pInsCx synapse (see discussion). The PPR was unchanged between the two groups (p = 0.91; Figure 5E, F). Thus, the MGm to pInsCx connection does not show a canonical LTP driven by the insertion of AMPA-receptors, nor does this connection show a presynaptic form of LTP which should have increased the PPR (Lucas et al., 2016).

We next performed measurements of the AMPA/NMDA ratio for the A1 → pInsCx synapse after fear learning (Figure 5H, I). Again, mice were assigned to a CS+US group (regular fear learning), or to a CS-only group (control; Figure 5J, N = 6 and 6 mice). Optogenetically-evoked EPSCs at this connection showed a significant increase of the absolute amplitudes of the AMPA-EPSCs, and of the AMPA/NMDA ratio in the CS+US group compared to the CS-only controls (p = 0.02 and 0.0008 respectively, Mann-Whitney test; Figure 5J, K; n = 48 cells in N = 6 mice and n = 46 cells in N = 6 mice for the CS+US and CS-only groups, respectively). On the other hand, the amplitude of the NMDA-EPSCs were unchanged (p = 0.94, Mann-Whitney test; Figure 5K), and the PPR was also unchanged between the two groups (p = 0.45, t-test; Figure 5M, N). These experiments show that the A1 → pInsCx synapse undergoes a postsynaptic form of LTP after fear learning, whereas at the MGm → pInsCx synaptic connection, fear learning does not induce a canonical form of LTP. Thus, two synaptic inputs that potentially carry auditory information to the pInsCx undergo differential long-term plasticity following fear learning.

### Transmission at the A1 → pInsCx synapse underlies *in-vivo* plasticity

We found that the A1 → pInsCx synapse undergoes a postsynaptic form of LTP after fear learning (Figure 5). It is possible, therefore, that LTP at this synapse drives the acquisition of CS-responses that we observed during fear learning (Figure 1). To test this idea, we performed *in-vivo* single-unit recordings, combined with optogenetic inhibition of afferent axons from the A1. Mice were injected with an AAV vector bilaterally in A1, to drive the expression of the inhibitory opsin Halorhodopsin (AAV8:hSyn:eNpHR3.0-eYFP), and an optrode was implanted in the left pInsCx (Figure 6A, B; Figure S3). Four weeks later, mice underwent the fear learning protocol, and *in-vivo* single-unit recordings were performed throughout all sessions. We applied short (300 ms) laser light pulses to silence the A1 axons during every second tone beep, on the habituation - and recall sessions (Figure 6C). This allowed us to compare the response to tone beeps in the presence- and absence of light in the same experiments, using the ‘no-light’ tone stimuli as internal controls.

**Figure 6.**
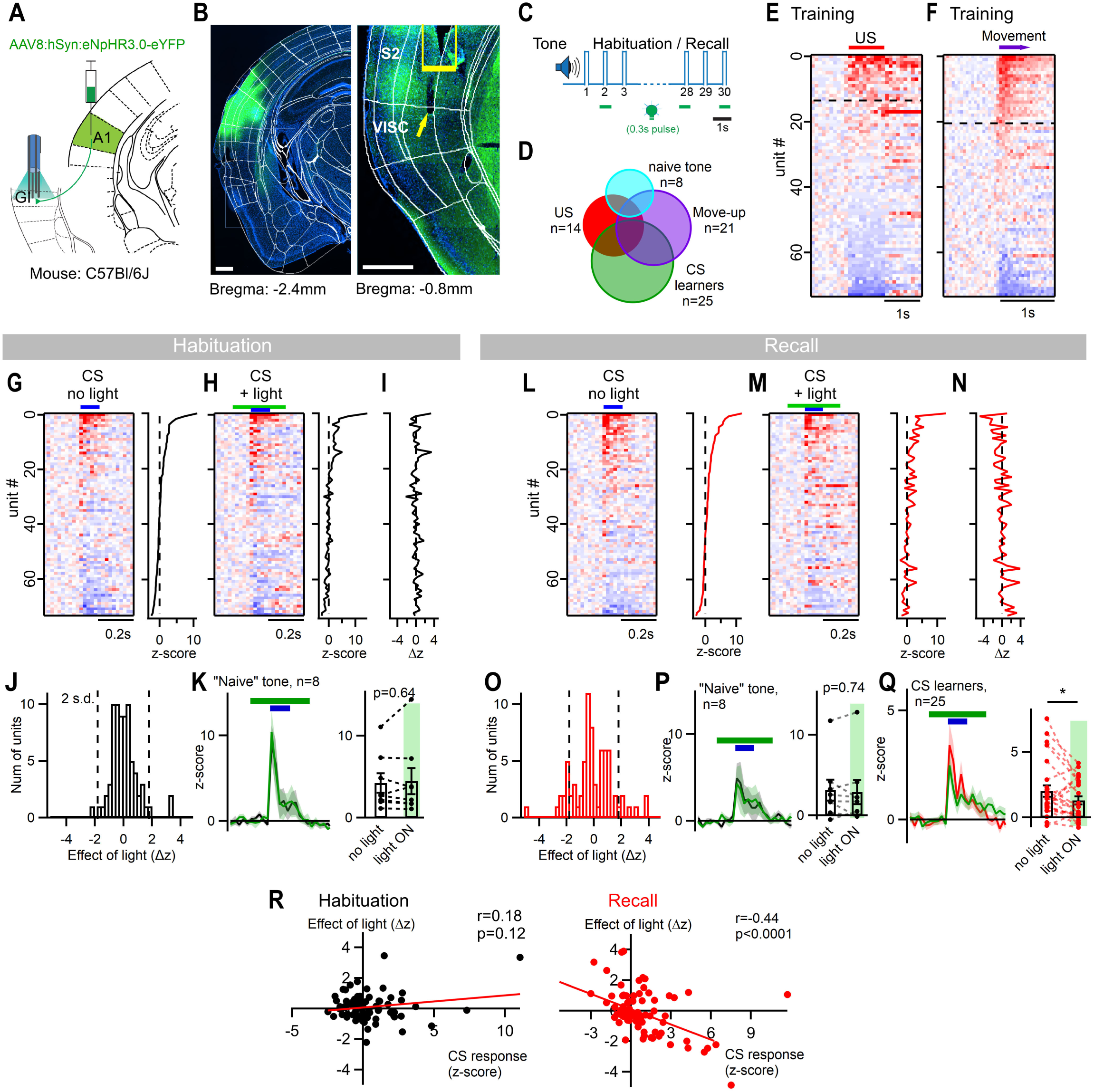
Functional synaptic transmission at the A1 → pInsCx connection is necessary to observe CS-learner responses during fear recall. ***A,*** Schematic of the experimental approach which combines optrode recordings in the pInsCx with optogenetic inhibition of incoming A1 axons. ***B,*** *Left*, image of the virus injection site in A1 (green channel, eNpHR3.0-eYFP; blue channel, DAPI). Right, image on the level of the pInsCx - S2 border with eNpHR3.0-eYFP expressing A1 axons (green channel) and the trace of a tetrode tip (arrow). The track of the optic fiber is marked by yellow box. ***C,*** Schematic of optogenetic axon silencing during every second tone beep during the habituation - and recall sessions. ***D,*** Venn diagram showing the sub-populations of pInsCx units with different response types in this experiment (N = 6 mice; n = 74 units). See also Figure 2J for comparison to the larger dataset in Figures 1 and 2. ***E,*** Raster plot of average aligned z-score responses to the footshock (US) presentation during the training session, sorted in decreasing order of response value (n = 74 units; N = 6 mice). Dashed line shows cut-off for positive US responders (n = 14). ***F,*** z-score responses aligned to the onset of movement (n = 21 movement-up responders, see dashed line). ***G,*** *Left*: Raster plot of aligned average responses to non-silenced tone beep presentations during the habituation session, sorted in decreasing order of response amplitude. *Right*: corresponding time-averaged z-score values observed during the tone beep. ***H,*** Display as in G, but shown for the tone beeps that received a 300 ms light pulse (see green line). The unit identities are matched to the ones in panel G. ***I,*** Differential z-score values (Δz) computed for each neuron as a difference in the z-score between optogenetically silenced CS beeps (H) and non-silenced control beeps (G). ***J,*** Histogram of Δz values from the data in panel I. ***K,*** *Left*: Tone responses of naïve CS responders during the habituation session, plotted separately for the non-silenced (black) and silenced beeps (green). Shading is ± s.e.m. *Right*: quantification of peak amplitudes of the tone responses for light, and no light conditions. ***L-M,*** Data shown as in panels G-I, here for the recall session. Note that the Δz plot reveals more negative values, and more scattering than during the habituation session (compare panels N and I). ***O***, Histogram of Δz values from the data in panel N. ***P,*** Data as in panel K, here shown for the recall session. As during the habituation session (see panel K), optogenetic inhibition of A1 axons had no effect on the naïve tone responders (n = 8, p = 0.74; Wilcoxon test). ***Q,*** *Left*: Tone responses of CS learners during the recall session, plotted for the non-silenced (red) and silenced tone beeps (green). Shading is ± s.e.m. *Right*: quantification of peak amplitudes of the tone responses for light, and no light conditions. Note the significant reduction of tone beep responses (n = 25, p = 0.038, paired t-test). ***R,*** Scatter plots of the modulation by light (Δz values) as a function of the tone beep response in the absence of light, both for the habituation session (*left*) and the recall session (*right*). Linear regression and Pearsońs correlation test revealed no significant correlation during habituation, but a significant negative correlation during the recall session (see Results for statistical parameters). Scale bars in B, 500 μm.

Similar as in the dataset of Figure 1, in these recordings we observed units that responded to footshocks (US; n = 14; Figure 6D, E), and to the movement onset after freezing (n = 21; Figure 6F; total number of n = 74 units recorded in N = 6 mice). Furthermore, we observed naive tone responders (n = 8) and CS-learners (n = 25, Figure 6D). During the habituation session, application of green light did not lead to a significant change of the tone response across the population of recorded units (Figure 6G-K, n = 8 naive tone responders, p = 0.64, paired t-test). For all recorded units, the distribution of the *difference* of the peak amplitude of the tone-driven AP-firing responses in the presence and absence of light pulses was symmetrical, indicating response variability (Figure 6I). Because tone responses during the habituation session are, by definition, naive tone responses, this data suggests that transmission at the A1 → pInsCx synapse is not necessary for this response type.

Interestingly, during the fear memory recall session, we observed a number of differences in the tone (CS) responses, and their modulation by yellow light (Figure 6L-Q). First, during the fear memory recall session, there was a larger number of units with tone responses (n = 33) than during the habituation session (n = 8). In agreement with the results in Figure 1, this suggests that previously silent CS-learner units have acquired a CS response after fear learning. Furthermore, during the recall session, green light application caused a significant decrease of the tone responses for the CS-learner units (Figure 6Q; p = 0.038; paired t-test), but not for the naive tone responders (Figure 6P; p = 0.74, Wilcoxon test). For all units recorded during the recall session, the light-induced *differences* of the peak response amplitudes showed a broader distribution as compared to the habituation day, with responses that were modulated in both the negative - and positive direction (compare Figure 6I, J for habituation, with Figure 6N, O for the recall session). Furthermore, there was a significant negative correlation between the modulating effect of light, and the amplitude of the CS-evoked AP-firing response during the recall session, but not during the habituation session (Figure 6R; recall session: r = -0.44, p < 0.0001; habituation session: r = 0.18, p = 0.12; Pearson’s test; n = 74 for both sessions). The *increased* amplitudes of CS-driven AP-firing in some units might indicate that optogenetic inhibition of afferent axons from A1 caused a reduction of the feedforward inhibition that dominated the response modulation; note that above we found evidence for feedforward inhibition from the A1 to the pInsCx (Figure 4). Thus, units with small or negative responses tended to be up-regulated by the optogenetic inhibition of A1 axons, whereas units with large CS-responses became down-regulated (Figure 6R). Taken together, optogenetic silencing of A1 axons in the pInsCx in combination with single-unit recordings reveals that *in-vivo*, transmission at the A1 → pInsCx synapse supports AP-firing of CS-learner units during fear memory recall; that is, after fear learning-induced plasticity has occurred. On the other hand, naive tone responses during the habituation session prior to fear learning were not affected by silencing of A1 axons. Together with our *ex-vivo* findings showing an increased AMPA/NMDA ratio at the A1 → pInsCx synapse (Figure 5), this data strongly suggests that potentiated synaptic transmission at the synaptic inputs from A1 causes the acquisition of a CS response in the pInsCx.

## Discussion

Using *in-vivo* single-unit recordings in the mouse model, we find a sub-population of neurons in the pInsCx which acquires a response to an auditory CS during fear learning. These neurons are enriched at the pInsCx - S2 border, in an area that corresponds to the IAF (Rodgers et al., 2008; Sawatari et al., 2011; Gogolla et al., 2014; Vestergaard et al., 2023; Jankowski et al., 2025). Anatomy and *ex-vivo* optogenetic circuit mapping show that the IAF receives excitatory, glutamatergic synapses from a non-lemniscal part of the auditory thalamus (MGm), and from the primary auditory cortex (A1). Measurements of AMPA/NMDA ratios at each optogenetically-stimulated connection revealed that the synapse from A1 → pInsCx, but not the one from MGm → pInsCx, undergoes a canonical form of LTP after fear learning. Furthermore, *in-vivo* single-unit recordings combined with optogenetic silencing of afferent axons showed that the acquired CS-response depends on unperturbed synaptic transmission at the A1 → pInsCx connection. Thus, our study shows that neurons in the pInsCx acquire a CS response via LTP of an associative cortical input from A1. This plastic CS representation is further integrated with a movement-up / freezing down representation in the pInsCx. Below, we discuss the implication of these findings for our understanding of the neural circuit mechanisms of fear learning.

### *In-vivo* activity of pInsCx neurons during fear learning

Our *in-vivo* recordings showed that a sub-population of neurons localized at the pInsCx - S2 border acquire a CS response during fear learning. Anterograde labelling from the auditory thalamus and from the A1 revealed spatially confined axon bundles at the same dorso-ventral position, and optogenetic circuit mapping showed that these axons generate glutamatergic EPSCs in neurons at the pInsCx - S2 border. Furthermore, we used a transgenic approach to label layer 4 neurons in sensory cortex, to validate that the auditory afferent axons are localized at the pInsCx - S2 border. Taken together, these data show that the CS-learner units are localized in the IAF, and that the IAF receives synaptic inputs from A1 and the MGm.

Beyond the responses to the auditory CS, the *in-vivo* recordings revealed further responses of pInsCx neurons during fear learning. First, ∼ 30% of the pInsCx neurons responded with an increase in AP-firing to the footshock stimulation employed as US in fear learning. In addition, there were neurons whose AP-firing activity was inhibited by the US, and yet other ones which responded after the termination of the US, called “US-offset” responders (Figure 1F, G; Palchaudhuri et al., 2025). Many “CS-learner” units showed a positive US response (n = 22/31 or 71%, Figure 2J). This finding supports evidence for models that posit that US-driven neuronal activity drives associative long-term plasticity at CS-encoding synapses, as it has been proposed previously for synapses in the lateral amygdala (Rumpel et al., 2005; Johansen et al., 2010; see Sigurdsson et al., 2007 and Palchaudhuri et al., 2022 for reviews).

Interestingly, we have also found robust coding of movements and freezing in the pInsCx. Thus, a subpopulation of the recorded units transiently increased their AP-firing when the mice initiated locomotion after freezing; during freezing these neurons tonically decreased their activity (movement-up / freezing-down responses, Figure 2A-H). A smaller subset of units showed the opposite regulation by movement, i.e. freeze-up / movement-down responses (Figure 2J). A previous *in-vivo* Ca^2+^ imaging study in head-fixed mice found freeze-up responses in a comparable subpopulation of neurons, but movement-up responses were not reported (Gehrlach et al., 2019). Another study, employing fiber photometry Ca^2+^ measurements of the pInsCx in freely moving mice during fear learning, reported a *decrease* in Ca^2+^ upon freezing onset (Klein et al., 2021); this response might reflect the activity of movement-up / freezing-down neurons which we describe here.

Several recent studies have described movement-related activity in sensory cortical areas of the mouse (Niell and Stryker, 2010; Keller et al., 2012; Schneider et al., 2014; Parker et al., 2020). Such movement-related coding in sensory areas has been implicated in the correction of perception errors caused by self-movement (Parker et al., 2020). The movement-up responses we describe here in the pInsCx are enhanced after fear learning and therefore, seem to undergo fear-learning related plasticity. It is reasonable to assume that in the context of fear learning, the freezing state entails a certain level of safety, by allowing the subject to avoid detection by predators in natural situations (Fanselow and Lester, 1988; Tseng et al., 2023). In turn, exploratory behaviors, maybe especially in the presence of a learned CS, implicate a less-safe condition in which enhanced alertness is indicated. Furthermore, our finding that significant movement-up responses occur only after immobility longer than ∼ one second (Figure 2D), is consistent with an aversive connotation of movement-onset coding.

Indeed, aversively-motivated freezing is likely longer than a spontaneous pause of movement that might last for less than a second. Thus, in one possible model, the movement-onset information in the pInsCx could be a preparatory signal for heightened alertness when animals break their freezing behavior and revert to exploration. Another interpretation would be that the transient movement-up responses are a pre-motor signal that contributes, possibly together with more classical motor areas, to drive movement initiation.

We have previously observed movement-up responses in direct- and indirect pathway neurons of the ventral tail striatum (vTS) during fear learning (Kintscher et al., 2023). Because the vTS receives strong excitatory inputs from the pInsCx (Hunnicutt et al., 2016; Kintscher et al., 2023) it is possible that the pInsCx transmits movement-up responses to the vTS. In addition, the vTS receives cortical inputs from auditory cortical areas, and therefore might be another site of integration of CS information, and movement-onset information. Further work is necessary, possibly including the investigation of upstream brain areas that carry movement information to the insula, to shed light on the functional implications of the robust movement-up representation that we have uncovered here in the pInsCx.

### Synaptic mechanisms of CS-plasticity in the insular cortex

We showed that the IAF of the pInsCx receives two synaptic inputs relevant for auditory coding: A projection from the MGm, and another one from the A1. Both inputs converge in the same dorso-ventral region of the IAF; they terminate in layer 4 in the case of the MGm and in layer 2/3 and -5 for inputs from the A1 (Figure 4D). In *ex-vivo* optogenetic stimulation experiments following fear learning, we showed that the A1 → pInsCx synapse undergoes a canonical form of LTP with increased AMPA-receptor mediated EPSCs, whereas the MGm → pInsCx synapse shows no increase in the AMPA/NMDA ratio. Nevertheless, the latter synapse shows an increased NMDA-EPSC; this plasticity might be similar to NMDA-plasticity described earlier at hippocampal mossy fiber synapses (Kwon and Castillo, 2008; Rebola et al., 2008). Thus, together with its location in layer 2/3 and -5 and its canonical form of LTP, the A1 → pInsCx connection can be regarded as an associative input, whereas the MGm → pInsCx synapse appears to be a more basic input potentially carrying less-processed auditory, and somatosensory information (Barsy et al., 2020; Taylor et al., 2021). Consistent with the findings of an increased AMPA/NMDA ratio at the A1 → pInsCx synapse, we found that unperturbed synaptic transmission at the A1 → pInsCx connection is necessary to observe the acquired CS-response during fear recall (Figure 6). This pinpoints the A1 → pInsCx connection as the site of plasticity that underlies the fear-learning induced acquisition of a CS in the pInsCx.

### Possible behavioral role of a fear-related CS-representation in the pInsCx

We have shown that the IAF of the pInsCx contains a significant fraction of neurons that acquire a CS response after fear learning (∼ 20% at the pInsCx - S2 border; ∼ 8% in the entire area of the pInsCx investigated here; Figure 1K). This fraction is similar to the one in the lateral amygdala of rats after fear learning (Quirk et al., 1995), and in the mouse basolateral amygdala (Grewe et al., 2017). The question then arises, do the acquired CS responses in the pInsCx contribute to the formation - and/or retrieval of an auditory-cued fear memory? To address this possibility, we have recently optogenetically silenced the activity of pInsCx principal neuron during the US (Palchaudhuri et al., 2025), and during the CS presentations during fear memory recall (Osypenko, 2023; Figure 3.14). In these experiments, we did not observe a significant reduction of CS-driven freezing during fear memory recall (Osypenko, 2023; Palchaudhuri et al., 2025). At face value, these experiments have not produced evidence for a role for CS-plasticity in the pInsCx for the formation, nor for the retrieval of an auditory-cued fear memory. Nevertheless, possible technical limitations need to be considered. First, it remains possible that limited tropism of AAV vectors for certain neuron subtypes (Aschauer et al., 2013; Scheyltjens et al., 2015; Challis et al., 2022) has not allowed us to target a sufficiently large number of pInsCx principal neurons, or else a relevant neuronal sub-population of the pInsCx, in our recent silencing experiments. Furthermore, fiber placements mainly target layer 5 of the pInsCx in our experiments (Figure 1K, L; (Osypenko, 2023; Palchaudhuri et al., 2025). Therefore, it remains possible that neuronal populations in other cortical layers relevant for CS plasticity, were not sufficiently covered by light illumination. Alternative methods of light application, like laterally implanted LED arrays that allow illumination through the entire cortical depth, might be advantageous for future optogenetic silencing experiments in the insular cortex (Takemoto et al., 2023).

Many studies including ours have employed freezing as a favorable measure of a conditioned response modulated by fear learning (Fanselow, 1982; LeDoux, 2000). Importantly, fear learning also goes along with physiological - and autonomic changes, like changes in blood pressure and heart rate (Kapp et al., 1979; LeDoux et al., 1988; Signoret-Genest et al., 2023). Given that the pInsCx is involved in visceral sensory processing and body - brain interactions (Cechetto and Saper, 1987; Yasui et al., 1991; Klein et al., 2021; Hsueh et al., 2023), it will be relevant to investigate whether the aversively-motivated auditory memory trace we found here in the pInsCx, might exert control over autonomic - or physiological responses in fear learning.

In summary, we have found that the IAF of the pInsCx elaborates a plastic CS-representation during auditory-cued fear learning, and we have unravelled the underlying synaptic mechanisms, which consist of LTP at an associative A1 → pInsCx connection. Future work should investigate the behavioral role(s) of a plastic representation of sensory events in the pInsCx during aversive forms of associative learning.

## Acknowledgements

We thank Heather Murray and Melanie Sipion for mouse genotyping and expert technical assistance. Fluorescence microscopy was performed at the BioImaging and Optics platform (BIOP) of EPFL. This research was supported by grants from the Swiss National Science Foundation (SNSF) to R.S. (31003A_176332 / 1 and 310030_204587 / 1).

## Methods

### Experimental animals

All experimental procedures with wildtype or genetically modified mice (*Mus musculus*) were approved by the veterinary office of the Canton of Vaud, Switzerland (authorizations VD3518 and VD3518×1). Mice of the C57Bl/6J strain (JAX stock #000664; RRID:IMSR_JAX:000664) were purchased from Charles River Laboratories (France). For some experiments, transgenic mouse lines were used: *Scnn1a^Cre^*(B6;C3-Tg(Scnn1a-cre)3Aibs/J; RRID:IMSR_JAX:009613; Madisen et al., 2010, *PV^Cre^*(B6;129P2*-Pvalb^tm1(cre)Arbr^*/J; RRID:IMSR_JAX:017320; Hippenmeyer et al., 2005, and a tdTomato reporter line, *Rosa26LSL-tdTomato* (B6.Cg-Gt(ROSA)26Sor^tm9(CAG-tdTomato)Hze^/J; RRID:IMSR_JAX:007909; also known as Ai9; Madisen et al., 2010. The latter was crossed with *Scnn1a^Cre^* or *PV^Cre^* lines for labelling of layer 4 neurons, or PV-interneurons, respectively. Transgenic mouse lines were back-crossed to a C57Bl/6J background for at least five generations.

Mice were group-housed under a 12/12 h light/dark cycle (lights on at 7am) with food and water *ad libitum*. One day prior to stereotaxic surgery, 6 - 7 weeks old male mice from 1-2 cages were randomly selected for experiments, after which they were housed in single cages until the end of experiment. We studied only male mice because fear learning shows sex-specific differences in rodents (Maren et al., 1994; Pryce et al., 1999; Gruene et al., 2015). Therefore, investigating both sexes in a non-differentiated way would have increased the variability of the behavioral results.

### Viral vectors and tracers

For anterograde labeling of axonal projections, we used AAV2/8:hSyn:eGFP (cat. 50465-AAV8, Addgene, Watertown, MA, USA) or AAV2/5:hSyn:eGFP (cat. v81-5, University of Zürich Viral Vector Facility, UZH-VVF, Zürich, Switzerland). For optogenetic circuit mapping experiments, AAV2/8:hSyn:Chronos-eGFP or AAV2/5:hSyn:Chronos-eGFP (University of North Carolina Vector Core, Chapel Hill, NC, USA) were injected into the A1 or the MGm, respectively. To drive expression of Halorhodopsin in A1 axons, AAV2/8:hSyn:eNpHR3.0-eYFP (cat. v560-8, UZH-VVF) was injected into A1. The volume of injected viral suspension was 200 nl. For retrograde labelling experiments, 50 nl of 0.5% Cholera toxin subunit B conjugated with Alexa Fluor 488 or Alexa Fluor 647 (CTB-488 or CTB-647 respectively; cat. C34775, C34778, Molecular Probes, OR, USA) were injected at two dorso-ventral positions of the pInsCx.

### Stereotaxic surgery

Stereotaxic surgery for virus or tracer injections and optrode implantation was performed in 6-7 weeks old male mice as previously described (Tang et al., 2020; Palchaudhuri et al., 2025). In brief, mice were anaesthetized with isoflurane (3% in O_2_ for induction and 1-1.5% in O_2_ for maintenance) and head-fixed using non-rupture ear-bars (Zygoma Ear cups, Kopf Instruments Model 921) in a stereotaxic frame (Model 940, David Kopf, Tujunga, CA, USA) on a heating pad (Harvard Apparatus, Holliston, MA, USA). Local anesthesia was achieved by subcutaneous injection of ∼50 μl of a mixture of lidocaine (1 mg/ml) and bupivacaine (1.25 mg/ml) in 0.9% saline. Craniotomies were made above the prospective injection or implantation sites. Viral vectors or CTB was injected using borosilicate glass micropipettes, at the following stereotaxic coordinates (anteroposterior, AP; mediolateral, ML; dorsoventral, DV; in mm): pInsCx, dorsal part, -0.9 AP, ±4.2 ML, -3.7 DV; pInsCx, ventral part, -0.9 AP, ±4.2 ML, -4.1 DV; MGm, -3 AP, ±2.1 ML, -3.25 DV; A1, -2.4 AP, ±4.3 ML, -2.4 DV.

A grounding - and an anchoring microscrews were inserted into two additional craniotomies; optrodes were slowly inserted to ∼600 μm above the desired coordinate of the recording tetrode tips, and were further adjusted by a microdrive 24h before the experiment. The skull surface and the microdrive were treated with light-curing adhesive (iBond Total Etch, Kulzer GmbH, Hanau, Germany) and embedded in the light-curing dental composite (Tetric EvoFlow, Ivoclar Vivadent, Schaan, Liechtenstein). After surgery, the animal was placed in its home cage under a heat lamp to recover under surveillance. One day prior and 6 days after the surgery, the drinking water was supplemented with 2 mg/ml paracetamol for analgesia.

### Auditory cued fear learning

Three weeks after surgery, the mice were habituated to handling and head tethering with the optrode cable and the fiber optic patch cord during five consecutive daily sessions (∼ 15 min each). On the following three days, the mice underwent auditory-cued fear learning, using a fear conditioning chamber (MED-VFC-OPTO-M, Med Associates Inc., Fairfax, VT, USA) placed inside a sound-attenuating enclosure (NIR-022MD, Med Associates Inc.). For all three sessions, a mouse at a time was placed in the conditioning chamber with optrode cables connected to its head. On day 1 (habituation session), six CS (tone) blocks were presented with a pseudo-random inter-CS block interval varying around 60s. Each tone block (30 s) consisted of 7 kHz, 80 dB tone beeps of 100 ms duration, repeated at 1 Hz. On day 2 (training session), six tone blocks (CS, as above), immediately followed by a 1s footshock (US; 0.6 mA AC, from a shock generator ENV-414S, Med Associates Inc) were presented. On day 3 (recall session), the mice were re-exposed to four tone blocks (4x CS). For the habituation- and training sessions, the fear conditioning chamber was configured for context A (rectangular plexiglass walls, metal grid floor, cleaned with 70% ethanol). For the recall session (day 3), the mice were placed in the same conditioning chamber, which was now configured for context B (semi-circular walls, smooth acrylic floor, cleaned with soap). A CMOS camera with an infrared filter continuously recorded the video of mice (30 frames/s).

For the measurements of AMPA/NMDA ratios after fear learning, the mice were randomly assigned to two groups. The first group underwent standard auditory fear conditioning as described above (“CS+US”). A control group underwent identical fear learning protocols, except that no footshocks were presented during the training session (“CS-only”).

Video recordings of mouse behavior were analyzed using the ezTrack package in Python (Pennington et al., 2019). The movement index trace from the ezTrack was further analyzed using custom procedures in IgorPro 7 (WaveMetrics Inc., Lake Oswego, OR, USA) as follows. If the movement index value fell below the threshold of 200 arbitrary units for at least 0.5 s, the “freezing” state was assigned to that time interval. The resulting binary trace of the freezing state was binned (10 s per bin) to compute the percent of time spent freezing, and then averaged across all mice in a given experimental group.

### In-vivo electrophysiology

The microdrive-mounted optrode devices were custom-made as previously described (Tang et al., 2020; Osypenko, 2023). One day before the start of the fear conditioning protocol, a mouse with an implanted optrode was anesthetized (i.p. injection of a mix containing ketamine and xylazine, 90 and 10 mg/kg body weight, respectively), and the optrode was advanced ventrally using the microdrive until a target depth of -3.7 to -4.1 mm DV was reached. During the fear learning protocol, *in-vivo* extracellular recordings were made continuously from the four tetrodes at 40 kHz sampling rate, using a 16-channel amplifier ME16-FAI-mPA under control of MC_Rack software (Multi Channel Systems, Reutlingen, Germany).

For combined *in-vivo* recordings with optogenetic inhibition of A1 axons, 300 ms green laser light pulses (561 nm) were applied during each second 100-ms tone beep during the habituation - and recall sessions. The light from a diode-pumped solid-state laser (MGL-FN-561-AOM-100mW; CNI Lasers, Changchun, China) was delivered via an optic fiber patch cord (200 μm core / 0.22 NA / 1 m; Doric Lenses Inc., Quebec, Canada). The power was adjusted to 10 mW at the optrode’s output with the laser’s acousto-optical modulator; residual power was blocked by a mechanical shutter during light-off times (SHB05T; Thorlabs Inc, Newton, NJ, USA).

### Analysis of in-vivo recordings

The raw data recorded by the MC_Rack software was converted to HDF5 format using Multi Channel Data Manager (MultiChannel Systems) and then processed (except the spike clustering step) using custom-written routines in IgorPro 7. The voltage traces were bandpass filtered (0.6–6 kHz for spike detection and 0.3-6 kHz for clustering) with a 4^th^ order Butterworth filter. The stimulation artifacts from electric footshocks (US, day 2) were semi-automatically blanked. Negative amplitude spikes were detected using a threshold (set at -3.8 SD for each electrode), and the timestamp of a spike was defined by the position of the largest negative peak across the channels of a given tetrode. Individual spike cutouts (spike timestamp ±1.6 ms) were clustered with the (KlustaKwik; (Rossant et al., 2016) algorithm using the MClust toolbox (Dr. David Redish; University of Minnesota) in MATLAB (MathWorks, Natick, MA, USA). Spike valley and the three principal components were used as clustering parameters. The clustering results were manually inspected, guided by the maximal value of the similarity metric: [RMS_i_ ∗ (1 − *ρρ*_*ii*_)]^−1^, and by overlapping the units waveforms. Here, RMS is the root mean square of a point-by-point difference between the tetrode-average waveforms of two putatively different units, and ρ_i_ is the average Pearson’s correlation coefficient calculated between the waveforms. Clusters with similar average waveforms which were not well separated in the valley-PC space, were likely from the same source and thus were merged into one. The quality of the resulting clusters was estimated by the isolation distance and L-ratio metrics (Schmitzer-Torbert et al., 2005) to control for Type I and Type II errors. Clusters with an isolation distance < 24 or L-ratio > 0.5 were excluded from the analysis (which reduced the total number of units ∼ twofold). Finally, we excluded units which had different spike amplitudes during the footshocks, or which spiked exclusively during footshocks, as those were likely electrical artifacts.

The longitudinal identity of individual units was matched across days semi-automatically, by first detecting the large values of the similarity metric (as above), followed by a manual validation step using waveform overlay. On average, ∼80% of the recorded units could be followed across all three days of the fear conditioning protocol. The experimenter remained blind to the activity pattern of the units until their longitudinal identity was established.

For the analysis of neuronal responses, z-scores and firing frequency (in Hz) for each unit were calculated by cumulating spikes into 20, 50 or 100 ms long time bins for the analysis of CS, movement/freezing and US responses, respectively. Z-scores of responses were calculated based on the local baseline taken as 1 s time interval before the onset of the US or movement, or as 0.8 s interval before the onset of the CS tone beeps.

The response types of the units were classified according to the following criteria. The unit was considered US-responsive if the time-averaged z-score during the stimulus (calculated from the average post-stimulus time histogram of six footshock) exceeded a value of 2, or if the z-score exceeded a value of 4 in at least one bin. For negative US responses, a z-score threshold of -2 was used. The CS-responses were calculated by averaging the z-score over the duration of the 100ms tone across all 30 beeps within a CS block. The CS responses were calculated for four times along the fear learning protocol: for the habituation session (using all six CS blocks), for the early training session (using CS blocks #1 and #2), for the late training session (CS blocks #5 and #6) and for the recall session (using all four CS blocks).

To classify the units based on the temporal evolution of their CS response, the above four values were used as features in hierarchical clustering performed on all putative principal neurons (using Euclidean distance metric and average linkage in the Agglomerative Clustering function of scikit-learn Python library). A unit was classified as movement-up or freeze-up responder if the average z-score during a 0.25s window following the respective inter-state transitions exceeded a value of 2.

For optrode recording combined with optogenetic inhibition, the baseline for z-score normalization of the CS responses was taken as the whole duration of the recording to avoid bias, in case optogenetic inhibition during a tone beep would have affected the local baseline of the next beep. Consequently, threshold values for responses were reduced 4-fold, to 0.5 for the 30xCS averaged z-scores, or to a value of 1 for the peak z-score responses to single CS beeps.

### Patch-clamp electrophysiology

Electrophysiological recordings in brain slices for optogenetic circuit mapping, and for optogenetic synapse stimulation after behavior (measurement of AMPA/NMDA ratio) were performed 4-5 weeks after stereotaxic surgery. A mouse at a time was euthanized by decapitation under isoflurane anesthesia (3% in O_2_), and the brain was rapidly extracted. For measurements of AMPA/NMDA ratios, the decapitation was done 15 - 30 minutes following the fear recall session. Coronal brain slices (300 μm) containing the pInsCx were made with a VT1000S vibratome (Leica Microsystems, Wetzlar, Germany) in ice-cold N-methyl-D-glutamine (NMDG) based solution (“slicing buffer“; Ting et al., 2014) containing (in mM): 110 NMDG, 2.5 KCl, 1.2 NaH_2_PO_4_, 20 HEPES, 25 Glucose, 5 Na-ascorbate, 2 Thiourea, 3 Na-pyruvate, 10 MgCl_2_, 0.5 CaCl_2_, pH 7.4, adjusted with HCl, and saturated with carbogen (95% O_2_ / 5% CO_2_). Brain slices were first stored for 7 min at 34°C in the slicing buffer, then transferred into a chamber with carbogen-saturated storage solution at room temperature, containing (in mM): 92 NaCl, 2.5 KCl, 30 NaHCO_3_, 1.2 NaH_2_PO_4_, 20 HEPES, 25 glucose, 5 Na-ascorbate, 3 Na-pyruvate, 2 MgCl_2_ and 2 CaCl_2_, pH 7.4.

Optogenetic circuit mapping experiments were done at room temperature (22-24°C) with a recording (extracellular) solution containing (in mM): 125 NaCl, 2.5 KCl, 25 NaHCO_3_, 1.2 NaH_2_PO_4_, 25 glucose, 0.4 Na-ascorbate, 1 MgCl_2_ and 2 CaCl_2_, pH 7.4 adjusted with NaOH and continuously bubbled with carbogen. Whole-cell patch-clamp recordings were done with an EPC-10/2 patch-clamp amplifier (HEKA Elektronik). The pipette solution contained (in mM): 145 K-gluconate, 6 KCl, 10 HEPES, 3 Na-phosphocreatine, 4 Mg-ATP, 0.3 Na_2_-GTP, 0.5 EGTA, adjusted to pH 7.2 with KOH. Neurons were visualized and recorded under an upright microscope (BX51WI; Olympus, Tokyo, Japan) equipped with a Dodt IR gradient contrast illumination, using a 60x / 0.9 NA water immersion objective (LUMPlanFl, Olympus) and a CMOS camera (C11440-52U; Hamamatsu Photonics K.K., Shizuoka, Japan) under control of μManager software (Edelstein et al., 2014). The fluorescence of tdTomato-positive neurons as well as excitation of Chronos were done with high-power 560 and 470 nm LEDs, respectively (CREE XP-E2; Cree Inc., Durham, NC, USA), custom-coupled into the epifluorescence port of the microscope. Chronos-expressing synaptic terminals were excited using 1 ms blue light pulses, and the resulting optically-evoked EPSCs were recorded at -70 mV. At least 10 consecutive sweeps were used to quantify EPSC amplitudes. To confirm that EPSCs were mediated by AMPA receptors, 10 μM of NBQX (2,3-dihydroxy-6-nitro-7-sulfamoyl-benzo[f]quinoxaline-2,3-dion) was added to the extracellular solution.

For the AMPA/NMDA ratio and PPR measurements, patch-clamp recordings were conducted at near-physiological temperature (34°C) maintained by a heated recording chamber RC-6GL/PM-1, an inline heater SHM-6 and a control unit TC-344B (Warner Instruments, Holliston, MA, USA). The pipette solution contained (in mM): 140 Cs-gluconate, 10 HEPES, 8 TEA-Cl, 5 Na-phosphocreatine, 4 Mg-ATP, 0.3 Na_2_-GTP, 5 EGTA, pH 7.2 adjusted with CsOH. The extracellular recording solution was supplemented with 5 μM gabazine (SR-95531) to exclude disynaptic IPSCs by blocking GABA_A_ receptors. The AMPA - and NMDA components of the EPSCs were recorded at -70 mV and at +50 mV, respectively. The PPR of the AMPA-EPSC at 50 ms interval was measured at -70 mV as a ratio of the 2^nd^ over the 1^st^ EPSC peak amplitudes.

All chemicals were from Sigma-Aldrich (Merck KGaA, Darmstadt, Germany), except HEPES, NaCl, KCl (Thermo Fisher Scientific, Waltham, MA, USA), MgCl_2_ (AppliChem, Darmstadt, Germany), gabazine (Abcam, Cambridge, UK) and NBQX (Tocris, Bio-Techne, Bristol, UK).

Patch-clamp recordings were analyzed using IgorPro 7 using custom scripts. AMPA-EPSCs were quantified as the peak inward current of the optogenetically-evoked EPSC measured at - 70 mV. The NMDA component of the EPSCs was estimated from the compound EPSC recorded at +50 mV, by subtracting the inverted AMPA-R component measured at -70 mV (the latter was scaled to match the rising slope of the compound EPSC).

### Histology and fluorescent imaging

For post-hoc histology, mice were deeply anaesthetized with i.p. injection of pentobarbital solution (150 mg/kg body weight), then transcardially perfused with 4% paraformaldehyde (PFA) in PBS. The extracted brains were post-fixed in 4% PFA overnight at 4°C then dehydrated in 30% sucrose PBS solution. The brains were cut into 40 μm coronal sections with a HM450 freezing microtome (Thermo Fisher Scientific, Waltham, MA, USA). Free-floating sections were mounted on Superfrost® Plus slides (Thermo Fisher Scientific), embedded in Fluoroshield DAPI mounting medium (Sigma-Aldrich) and imaged at the Bioimaging and Optics Platform of EPFL (BIOP) using a slide scanner VS120-L100 (Olympus) equipped with a 10x/0.4 NA objective. The images of brain sections were imported into QuPath (Bankhead et al., 2017), and semi-automatically registered to the reference brain atlas (Allen Brain Atlas, 2014) using the ABBA plugin for FIJI (Chiaruttini et al., 2022).

### Experimental Design and Statistical Analyses

No *a priori* sample size calculation was performed. Statistical analysis was performed in Prism (GraphPad, San Diego, CA, USA). Before choosing the statistical test, the distribution of the data was tested for normality using a Shapiro-Wilk test. If normality was confirmed, a paired or unpaired version of the two-tailed Student’s t-tests was used for two-sample data, as indicated. Otherwise, non-parametric two-tailed tests were used: Wilcoxon matched pairs signed-rank test for paired comparisons, or Mann-Whitney U test for unpaired comparisons, as indicated. For statistical comparison of fractions of responding units, Fisher’s exact test was used. For comparison of multiple fractions, the p-values from individual Fisher’s exact tests were adjusted with a Bonferroni correction. For datasets with multiple factors and repeated measurements, a repeated measures ANOVA was used, followed by Tukey’s or Dunnett’s post-hoc tests as indicated. In case the data was non-normally distributed, a Friedman test followed by Dunn’s post-hoc test was used.

Each specific statistical test along with the sample size, values of the statistics and the p-values is mentioned in the Results text. The averaged data are expressed as mean ± SEM. Statistical significance, if applicable, is indicated in the Figures using asterisks as p ≤ 0.05 (*), p ≤ 0.01 (**) and p ≤ 0.001 (***).

## Supplementary Figure Legends

**Figure S1.**
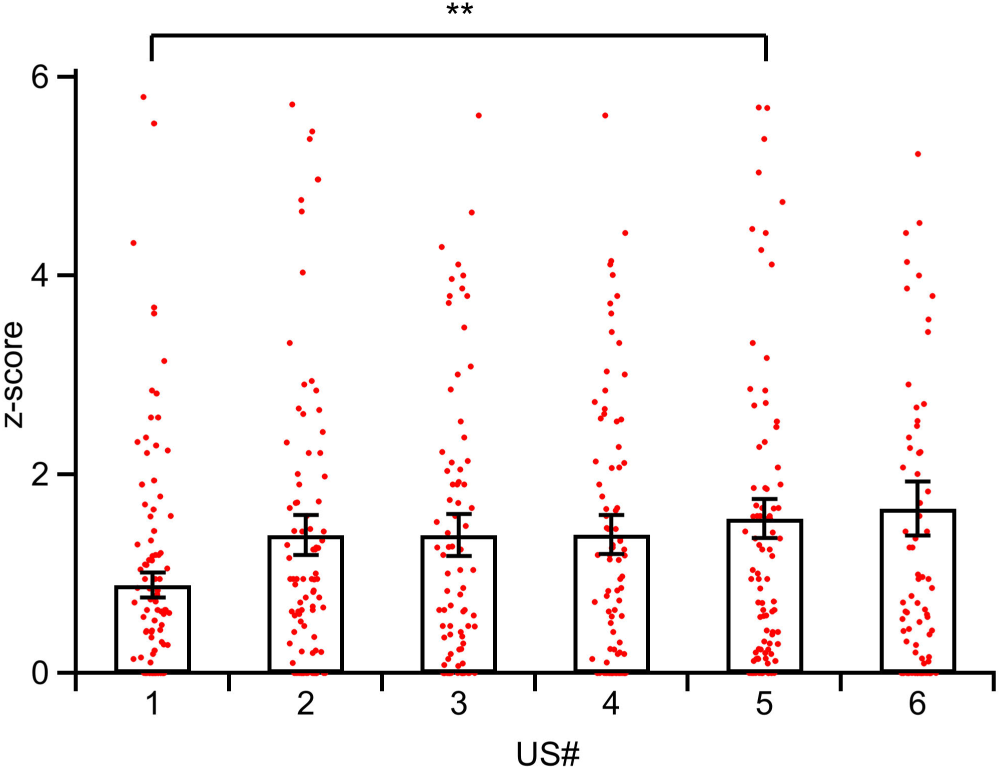
Dependence of footshock-response amplitude on footshock number upon repeated US presentation during the training session. The footshock-evoked AP-firing response amplitude (z-score) is shown for individual units (markers) and as average ± SEM values (bars), for the six subsequent US applications during the training session (n = 134 positive US-onset responders out of n = 374 putative principal neurons; see Figure 1F). Friedman test found an effect of footshock number (p = 0.011). A Dunn’s multiple comparison test reported a significantly smaller US-response to footshock #1, as compared to footshock #5 (p = 0.0049); all other comparisons were not significant.

**Figure S2.**
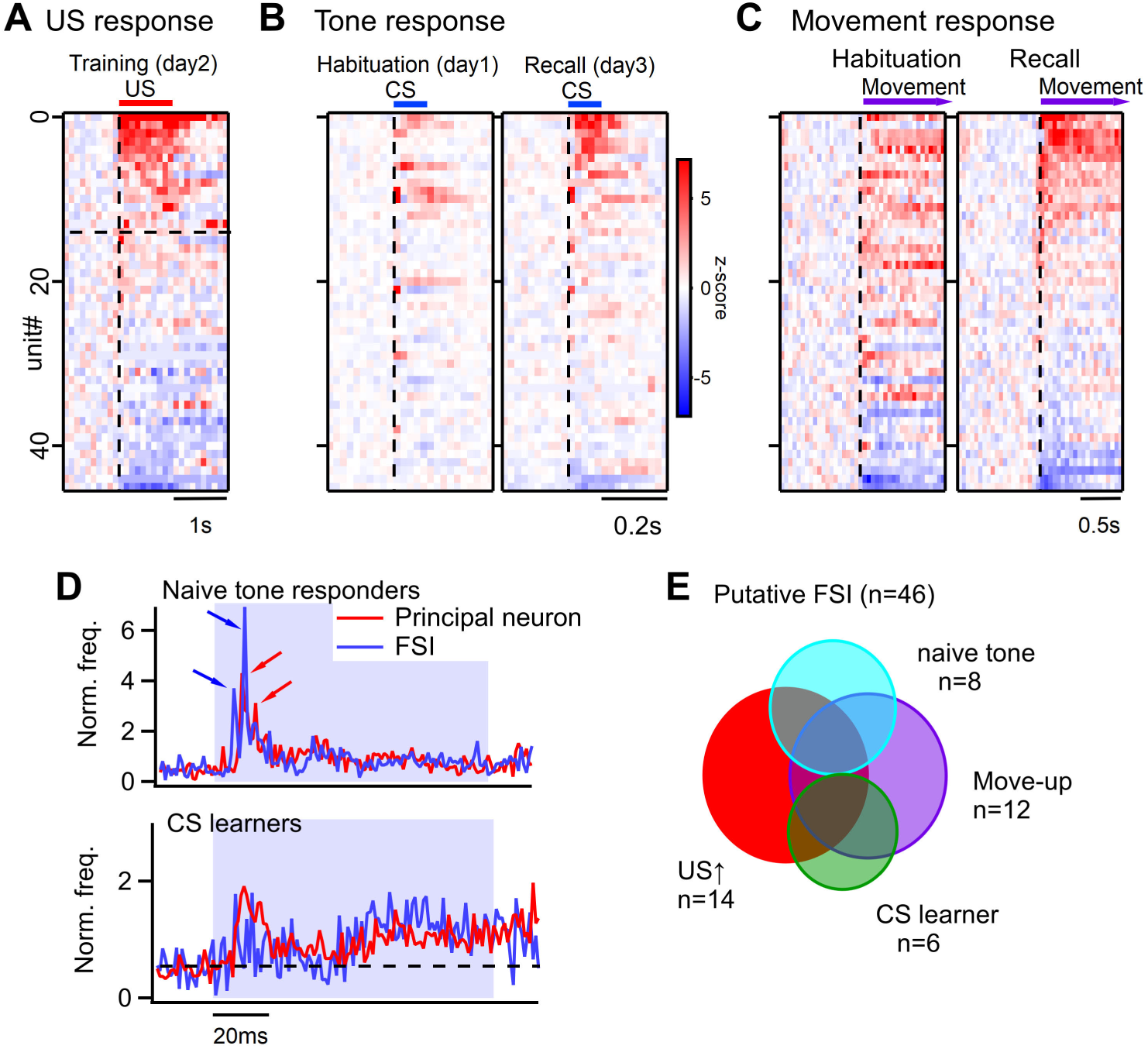
Responses of units classified as putative FSIs (fast spiking interneurons) during auditory fear learning. The figure summarizes responses of the units that were classified as fast spiking interneurons (FSI) based on their fast waveform characteristics (see Figure 1D; peak-to-trough waveform < 0.35 ms; n = 46 units). ***A,*** Raster plot for the averaged (n = 6) US-onset responses (see also Figure 1F for the putative principal neuron population). n = 14 units passed the threshold for classification of US-excited FSIs (see horizontal dashed line). ***B,*** Raster plots showing the average PSTH of responses to individual tone beeps during the habituation (*left*) - and recall session (*right*), sorted in decreasing order of the response amplitude during the recall session. Note that, similar to putative principal neurons (Figure 1I), some FSIs showed naïve tone responses. ***C,*** Average movement-up responses, aligned to the onset of the movement in each animal, both for the habituation session (left) and for the recall session (right). Compare to Figure 2F for the corresponding data for putative principal neurons. ***D,*** Baseline-normalized, time-resolved AP-firing frequencies for naive tone responders (*top*; both FSIs [blue trace, n = 8] and principal neurons [red trace, n = 12]), and for tone learners (*bottom*), in response to the 100 ms tone beeps. Blue shading shows the time of tone beep presentation. Note the two distinct short-latency peaks in the response of naïve tone responding FSIs (blue arrows) at 7 and 11 ms, respectively, after the tone onset. The response of naïve tone responding principal neurons also shows such peaks at somewhat longer latencies of 10 and 15 ms, respectively (red arrows). These sharp peaks might be driven by auditory inputs from the auditory thalamus (see also Jankowski et al., 2025 for short-latency auditory responses in the IAF). Interestingly, CS learners do not show such peaks. ***E,*** Venn diagram summarizing the overlap of sub-populations of n = 46 putative FSIs showing US-, movement-up, CS learner and naive tone response characteristics. Compare to Figure 1J for the corresponding analysis for putative principal neurons.

**Figure S3.**
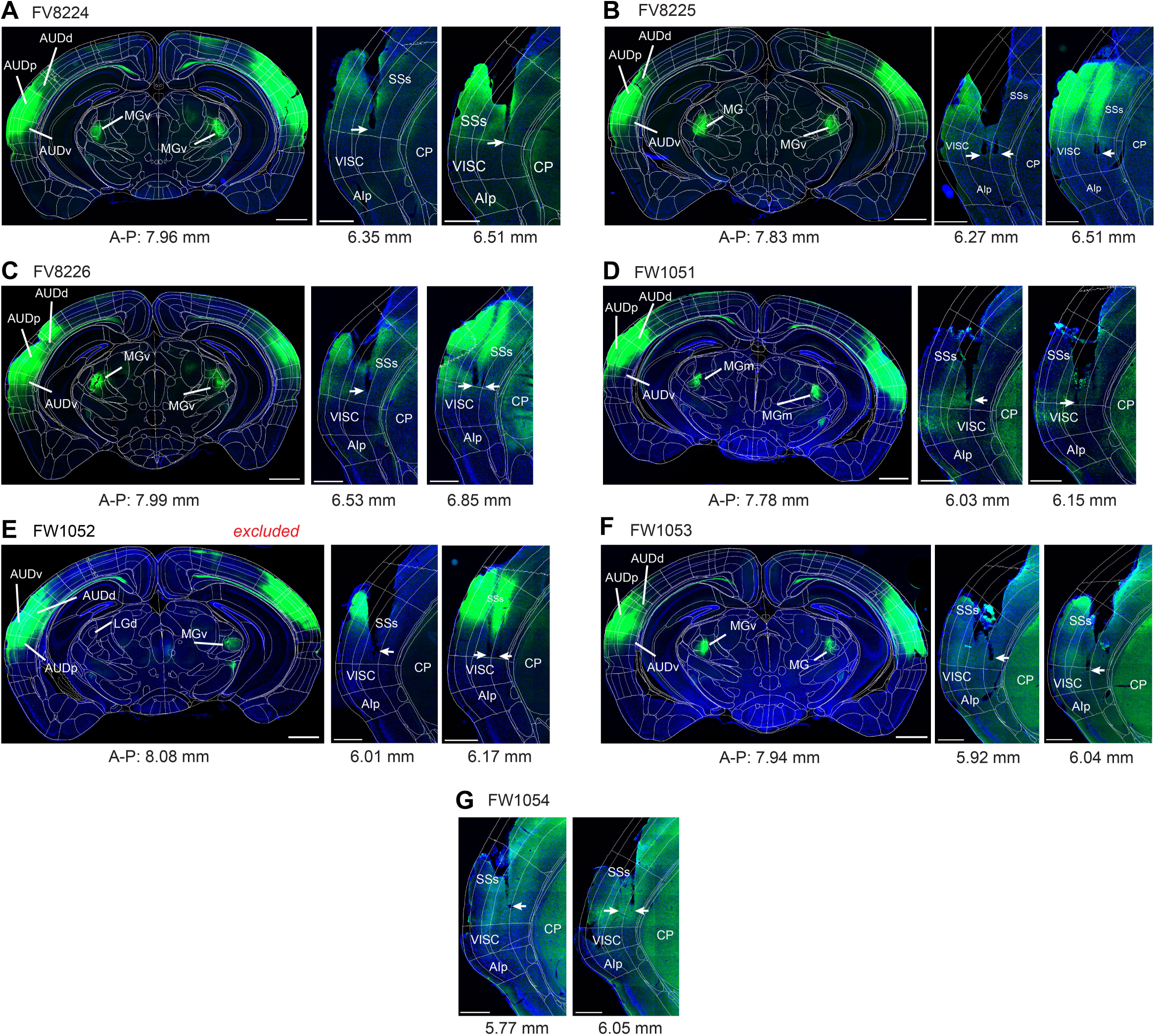
Histological validation of virus injection and optrode placement in the experiments of Figure 6. Images of coronal sections show the fluorescence of eYFP (green channel) resulting from the bilateral injection of AAV8:hSyn:eNpHR3.0-eYFP vector in the A1 (*left* panels), as well as the eYFP-positive axons and the tetrode tips (white arrows) on the level of the pInsCx (*right* panels). Blue channel, DAPI. The coronal sections were registered to the digital Allen Brain Atlas (ABA) using the QuPath-ABBA workflow (see Methods); the A-P numbers indicate antero-posterior location in the ABA coordinate system. Note the eYFP signals in the auditory thalamus (labelled MGv) in all mice except one (panel E, FW 1052). Cortico-thalamic outputs to the auditory thalamus should indicate a successful injection into the auditory cortex. Correspondingly, mouse FW1052 which unilaterally did not exhibit eYFP fluorescence in the auditory thalamus, was excluded from the dataset. For mouse FW1054, we failed to obtain coronal sections containing A1 (panel G).

## References

Allen Brain Atlas (2014). Allen Institute for Brain Science. Allen Mouse Brain Atlas. [Internet]. Available at http://mouse.brain-map.org/.

Aschauer, D.F., Kreuz, S., and Rumpel, S. (2013). Analysis of transduction efficiency, tropism and axonal transport of AAV serotypes 1, 2, 5, 6, 8 and 9 in the mouse brain. PloS one 8, e76310.

Bankhead, P., Loughrey, M.B., Fernandez, J.A., Dombrowski, Y., McArt, D.G., Dunne, P.D., McQuaid, S., Gray, R.T., Murray, L.J., Coleman, H.G., et al. (2017). QuPath: Open source software for digital pathology image analysis. Sci Rep 7, 16878.

Barsy, B., Kocsis, K., Magyar, A., Babiczky, A., Szabó, M., Veres, J.M., Hillier, D., Ulbert, I., Yizhar, O., and Mátyás, F. (2020). Associative and plastic thalamic signaling to the lateral amygdala controls fear behavior. Nature neuroscience 23, 625–637.

Bokiniec, P., Whitmire, C.J., Leva, T.M., and Poulet, J.F.A. (2022). Brain-wide connectivity map of mouse thermosensory cortices. Cerebral cortex.

Cechetto, D.F., and Saper, C.B. (1987). Evidence for a viscerotopic sensory representation in the cortex and thalamus in the rat. The Journal of comparative neurology 262, 27–45.

Challis, R.C., Ravindra Kumar, S., Chen, X., Goertsen, D., Coughlin, G.M., Hori, A.M., Chuapoco, M.R., Otis, T.S., Miles, T.F., and Gradinaru, V. (2022). Adeno-Associated Virus Toolkit to Target Diverse Brain Cells. Annual review of neuroscience 45, 447–469.

Chen, X., Gabitto, M., Peng, Y., Ryba, N.J., and Zuker, C.S. (2011). A gustotopic map of taste qualities in the mammalian brain. Science 333, 1262–1266.

Chiaruttini, N., Burri, O., Haub, P., Guiet, R., Sordet-Dessimoz, J., and Seitz, A. (2022). An Open-Source Whole Slide Image Registration Workflow at Cellular Precision Using Fiji, QuPath and Elastix. Front Comp Sci-Switz 3.

Coghill, R.C. (2020). The distributed nociceptive system: A framework for understanding pain. Trends in neurosciences 43, 780–794.

Dalmay, T., Abs, E., Poorthuis, R.B., Hartung, J., Pu, D.L., Onasch, S., Lozano, Y.R., Signoret-Genest, J., Tovote, P., Gjorgjieva, J., and Letzkus, J.J. (2019). A critical role for neocortical processing of threat memory. Neuron 104, 1180–1194 e1187.

Edelstein, A.D., Tsuchida, M.A., Amodaj, N., Pinkard, H., Vale, R.D., and Stuurman, N. (2014). Advanced methods of microscope control using muManager software. Journal of biological methods 1.

Fanselow, M.S. (1982). The postshock activity burst. Anim Learn Behav 10, 448–454.

Fanselow, M.S. (2018). The Role of Learning in Threat Imminence and Defensive Behaviors. Curr Opin Behav Sci 24, 44–49.

Fanselow, M.S., and Lester, L.S. (1988). A functional behavioristic approach to aversively motivated behavior: Predatory imminence as a determinant of the topography of defensive behavior. In: Evolution and Learning eds Bolles, R. C. & Beecher, M. D., 185–212.

Feinberg, T.E., and Mallatt, J.M. (2017). The ancient origins of consciousness: How the brain created experience. MIT Press.

Frontera, J.L., Baba Aissa, H., Sala, R.W., Mailhes-Hamon, C., Georgescu, I.A., Lena, C., and Popa, D. (2020). Bidirectional control of fear memories by cerebellar neurons projecting to the ventrolateral periaqueductal grey. Nature communications 11, 5207.

Gehrlach, D.A., Dolensek, N., Klein, A.S., Roy Chowdhury, R., Matthys, A., Junghänel, M., Gaitanos, T.N., Podgornik, A., Black, T.D., Reddy Vaka, N., et al. (2019). Aversive state processing in the posterior insular cortex. Nature neuroscience 22, 1424–1437.

Gogolla, N. (2017). The insular cortex. Curr Biol 27, R580–R586.

Gogolla, N., Takesian, A.E., Feng, G., Fagiolini, M., and Hensch, T.K. (2014). Sensory integration in mouse insular cortex reflects GABA circuit maturation. Neuron 83, 894–905.

Grewe, B.F., Gründemann, J., Kitch, L.J., Lecoq, J.A., Parker, J.G., Marshall, J.D., Larkin, M.C., Jercog, P.E., Grenier, F., Li, J.Z., et al. (2017). Neural ensemble dynamics underlying a long-term associative memory. Nature 543, 670–675.

Gruene, T.M., Flick, K., Stefano, A., Shea, S.D., and Shansky, R.M. (2015). Sexually divergent expression of active and passive conditioned fear responses in rats. eLife 4, e11352.

Herry, C., and Johansen, J.P. (2014). Encoding of fear learning and memory in distributed neuronal circuits. Nature neuroscience 17, 1644–1654.

Hippenmeyer, S., Vrieseling, E., Sigrist, M., Portmann, T., Laengle, C., Ladle, D.R., and Arber, S. (2005). A developmental switch in the response of DRG neurons to ETS transcription factor signaling. PLoS Biol 3, e159.

Hsueh, B., Chen, R., Jo, Y., Tang, D., Raffiee, M., Kim, Y.S., Inoue, M., Randles, S., Ramakrishnan, C., Patel, S., et al. (2023). Cardiogenic control of affective behavioural state. Nature 615, 292–299.

Hu, H., Gan, J., and Jonas, P. (2014). Interneurons. Fast-spiking, parvalbumin(+) GABAergic interneurons: from cellular design to microcircuit function. Science 345, 1255263.

Hunnicutt, B.J., Jongbloets, B.C., Birdsong, W.T., Gertz, K.J., Zhong, H., and Mao, T. (2016). A comprehensive excitatory input map of the striatum reveals novel functional organization. eLife 5.

Janak, P.H., and Tye, K.M. (2015). From circuits to behaviour in the amygdala. Nature 517, 284–292.

Jankowski, M.M., Karayanni, M., Harpaz, M., Polterovich, A., and Nelken, I. (2025). A Rapid Anterior Auditory Processing Stream through the Insulo-parietal Auditory Field in the Rat. The Journal of neuroscience : the official journal of the Society for Neuroscience 45.

Johansen, J.P., Hamanaka, H., Monfils, M.H., Behnia, R., Deisseroth, K., Blair, H.T., and LeDoux, J.E. (2010). Optical activation of lateral amygdala pyramidal cells instructs associative fear learning. Proceedings of the National Academy of Sciences of the United States of America 107, 12692–12697.

Kapp, B.S., Frysinger, R.C., Gallagher, M., and Haselton, J.R. (1979). Amygdala central nucleus lesions: effect on heart rate conditioning in the rabbit. Physiol Behav 23, 1109–1117.

Keller, G.B., Bonhoeffer, T., and Hübener, M. (2012). Sensorimotor mismatch signals in primary visual cortex of the behaving mouse. Neuron 74, 809–815.

Kim, W.B., and Cho, J.H. (2017). Encoding of discriminative fear memory by input-specific LTP in the amygdala. Neuron 95, 1129–1146 e1125.

Kintscher, M., Kochubey, O., and Schneggenburger, R. (2023). A striatal circuit balances learned fear in the presence and absence of sensory cues. eLife 12.

Klein, A.S., Dolensek, N., Weiand, C., and Gogolla, N. (2021). Fear balance is maintained by bodily feedback to the insular cortex in mice. Science 374, 1010–1015.

Kusumoto-Yoshida, I., Liu, H., Chen, B.T., Fontanini, A., and Bonci, A. (2015). Central role for the insular cortex in mediating conditioned responses to anticipatory cues. Proceedings of the National Academy of Sciences of the United States of America 112, 1190–1195.

Kwon, H.B., and Castillo, P.E. (2008). Long-term potentiation selectively expressed by NMDA receptors at hippocampal mossy fiber synapses. Neuron 57, 108–120.

LeDoux, J.E. (2000). Emotion circuits in the brain. Annual review of neuroscience 23, 155–184.

LeDoux, J.E., Iwata, J., Cicchetti, P., and Reis, D.J. (1988). Different projections of the central amygdaloid nucleus mediate autonomic and behavioral correlates of conditioned fear. The Journal of neuroscience : the official journal of the Society for Neuroscience 8, 2517–2529.

Little, J.P., and Carter, A.G. (2012). Subcellular synaptic connectivity of layer 2 pyramidal neurons in the medial prefrontal cortex. The Journal of Neuroscience 32, 12808–12819.

Lucas, E.K., Jegarl, A.M., Morishita, H., and Clem, R.L. (2016). Multimodal and site-specific plasticity of amygdala parvalbumin interneurons after fear learning. Neuron 91, 629–643.

Madisen, L., Zwingman, T.A., Sunkin, S.M., Oh, S.W., Zariwala, H.A., Gu, H., Ng, L.L., Palmiter, R.D., Hawrylycz, M.J., Jones, A.R., et al. (2010). A robust and high-throughput Cre reporting and characterization system for the whole mouse brain. Nature neuroscience 13, 133–140.

Maffei, A., Haley, M., and Fontanini, A. (2012). Neural processing of gustatory information in insular circuits. Current opinion in neurobiology 22, 709–716.

Maren, S. (2001). Neurobiology of Pavlovian fear conditioning. Annual review of neuroscience 24, 897–931.

Maren, S., De Oca, B., and Fanselow, M.S. (1994). Sex differences in hippocampal long-term potentiation (LTP) and Pavlovian fear conditioning in rats: positive correlation between LTP and contextual learning. Brain Res 661, 25–34.

McKernan, M.G., and Shinnick-Gallagher, P. (1997). Fear conditioning induces a lasting potentiation of synaptic currents in vitro. Nature 390, 607–611.

Mombelli, E., Osypenko, D., Palchaudhuri, S., Sourmpis, C., Brea, J., Kochubey, O., and Schneggenburger, R. (2024). Auditory stimuli suppress contextual fear responses in safety learning independent of a possible safety meaning. Front Behav Neurosci 18, 1415047.

Nicolas, C., Ju, A., Wu, Y., Eldirdiri, H., Delcasso, S., Couderc, Y., Fornari, C., Mitra, A., Supiot, L., Vérite, A., et al. (2023). Linking emotional valence and anxiety in a mouse insula-amygdala circuit. Nature communications 14, 5073.

Niell, C.M., and Stryker, M.P. (2010). Modulation of visual responses by behavioral state in mouse visual cortex. Neuron 65, 472–479.

Nieuwenhuys, R. (2012). The insular cortex: a review. Progress in brain research 195, 123–163.

Ogawa, H., Ito, S., Murayama, N., and Hasegawa, K. (1990). Taste area in granular and dysgranular insular cortices in the rat identified by stimulation of the entire oral cavity. Neuroscience research 9, 196–201.

Osypenko, D. (2023). Plasticity of sensory representations in the posterior insular cortex during fear learning. PhD Thesis n° 9335, Ecole Poleytechnique Fédérale de Lausanne EPFL.

Palchaudhuri, S., Lin, B.X., Osypenko, D., Wu, J., Kochubey, O., and Schneggenburger, R. (2025). A posterior insula to lateral amygdala pathway transmits US-offset information with a limited role in fear learning. Cell reports 44, 115320.

Palchaudhuri, S., Osypenko, D., and Schneggenburger, R. (2022). Fear learning: An evolving picture for plasticity at synaptic afferents to the amygdala. Neuroscientist, 10738584221108083.

Parker, P.R.L., Brown, M.A., Smear, M.C., and Niell, C.M. (2020). Movement-Related Signals in Sensory Areas: Roles in Natural Behavior. Trends in neurosciences 43, 581–595.

Pennington, Z.T., Dong, Z., Feng, Y., Vetere, L.M., Page-Harley, L., Shuman, T., and Cai, D.J. (2019). ezTrack: An open-source video analysis pipeline for the investigation of animal behavior. Sci Rep 9, 19979.

Petreanu, L., Huber, D., Sobczyk, A., and Svoboda, K. (2007). Channelrhodopsin-2-assisted circuit mapping of long-range callosal projections. Nature neuroscience 10, 663–668.

Pryce, C.R., Lehmann, J., and Feldon, J. (1999). Effect of sex on fear conditioning is similar for context and discrete CS in Wistar, Lewis and Fischer rat strains. Pharmacol Biochem Behav 64, 753–759.

Quirk, G.J., Repa, C., and LeDoux, J.E. (1995). Fear conditioning enhances short-latency auditory responses of lateral amygdala neurons: parallel recordings in the freely behaving rat. Neuron 15, 1029–1039.

Rebola, N., Lujan, R., Cunha, R.A., and Mulle, C. (2008). Adenosine A2A receptors are essential for long-term potentiation of NMDA-EPSCs at hippocampal mossy fiber synapses. Neuron 57, 121–134.

Ren, L., Fan, Y., Wu, W., Qian, Y., He, M., Li, X., Wang, Y., Yang, Y., Wen, X., Zhang, R., et al. (2024). Anxiety disorders: Treatments, models, and circuitry mechanisms. Eur J Pharmacol 983, 176994.

Ressler, K.J., Berretta, S., Bolshakov, V.Y., Rosso, I.M., Meloni, E.G., Rauch, S.L., and Carlezon, W.A., Jr. (2022). Post-traumatic stress disorder: clinical and translational neuroscience from cells to circuits. Nat Rev Neurol 18, 273–288.

Rich, M.T., Huang, Y.H., and Torregrossa, M.M. (2019). Plasticity at thalamo-amygdala synapses regulates cocaine-cue memory formation and extinction. Cell Rep 26, 1010–1020 e1015.

Rodgers, K.M., Benison, A.M., Klein, A., and Barth, D.S. (2008). Auditory, somatosensory, and multisensory insular cortex in the rat. Cerebral cortex 18, 2941–2951.

Rossant, C., Kadir, S.N., Goodman, D.F.M., Schulman, J., Hunter, M.L.D., Saleem, A.B., Grosmark, A., Belluscio, M., Denfield, G.H., Ecker, A.S., et al. (2016). Spike sorting for large, dense electrode arrays. Nature neuroscience 19, 634–641.

Roy, D.S., Park, Y.G., Kim, M.E., Zhang, Y., Ogawa, S.K., DiNapoli, N., Gu, X., Cho, J.H., Choi, H., Kamentsky, L., et al. (2022). Brain-wide mapping reveals that engrams for a single memory are distributed across multiple brain regions. Nature communications 13, 1799.

Rumpel, S., LeDoux, J., Zador, A., and Malinow, R. (2005). Postsynaptic receptor trafficking underlying a form of associative learning. Science 308, 83–88.

Sawatari, H., Tanaka, Y., Takemoto, M., Nishimura, M., Hasegawa, K., Saitoh, K., and Song, W.J. (2011). Identification and characterization of an insular auditory field in mice. The European journal of neuroscience 34, 1944–1952.

Scheyltjens, I., Laramée, M.E., Van den Haute, C., Gijsbers, R., Debyser, Z., Baekelandt, V., Vreysen, S., and Arckens, L. (2015). Evaluation of the expression pattern of rAAV2/1, 2/5, 2/7, 2/8, and 2/9 serotypes with different promoters in the mouse visual cortex. The Journal of comparative neurology 523, 2019–2042.

Schmitzer-Torbert, N., Jackson, J., Henze, D., Harris, K., and Redish, A.D. (2005). Quantitative measures of cluster quality for use in extracellular recordings. Neuroscience 131, 1–11.

Schneider, D.M., Nelson, A., and Mooney, R. (2014). A synaptic and circuit basis for corollary discharge in the auditory cortex. Nature 513, 189–194.

Signoret-Genest, J., Schukraft, N., Reis, S.L., Segebarth, D., Deisseroth, K., and Tovote, P. (2023). Integrated cardio-behavioral responses to threat define defensive states. Nature neuroscience 26, 447–457.

Sigurdsson, T., Doyere, V., Cain, C.K., and LeDoux, J.E. (2007). Long-term potentiation in the amygdala: a cellular mechanism of fear learning and memory. Neuropharmacology 52, 215–227.

Szönyi, A., Zichó, K., Barth, A.M., Gönczi, R.T., Schlingloff, D., Török, B., Sipos, E., Major, A., Bardóczi, Z., Sos, K.E., et al. (2019). Median raphe controls acquisition of negative experience in the mouse. Science 366.

Takemoto, M., Hasegawa, K., Nishimura, M., and Song, W.J. (2014). The insular auditory field receives input from the lemniscal subdivision of the auditory thalamus in mice. The Journal of comparative neurology 522, 1373–1389.

Takemoto, M., Kato, S., Kobayashi, K., and Song, W.J. (2023). Dissection of insular cortex layer 5 reveals two sublayers with opposing modulatory roles in appetitive drinking behavior. iScience 26, 106985.

Tang, W., Kochubey, O., Kintscher, M., and Schneggenburger, R. (2020). A VTA to basal amygdala dopamine projection contributes to signal salient somatosensory events during fear learning. The Journal of neuroscience : the official journal of the Society for Neuroscience 40, 3969–3980.

Taylor, J.A., Hasegawa, M., Benoit, C.M., Freire, J.A., Theodore, M., Ganea, D.A., Innocenti, S.M., Lu, T., and Gründemann, J. (2021). Single cell plasticity and population coding stability in auditory thalamus upon associative learning. Nature communications 12, 2438.

Ting, J.T., Daigle, T.L., Chen, Q., and Feng, G. (2014). Acute brain slice methods for adult and aging animals: application of targeted patch clamp analysis and optogenetics. Methods in molecular biology 1183, 221–242.

Tovote, P., Fadok, J.P., and Lüthi, A. (2015). Neuronal circuits for fear and anxiety. Nature reviews. Neuroscience 16, 317–331.

Tseng, Y.T., Schaefke, B., Wei, P., and Wang, L. (2023). Defensive responses: behaviour, the brain and the body. Nature reviews. Neuroscience 24, 655–671.

Vestergaard, M., Carta, M., Guney, G., and Poulet, J.F.A. (2023). The cellular coding of temperature in the mammalian cortex. Nature.

Wang, Q., Zhu, J.J., Wang, L., Kan, Y.P., Liu, Y.M., Wu, Y.J., Gu, X., Yi, X., Lin, Z.J., Wang, Q., et al. (2022). Insular cortical circuits as an executive gateway to decipher threat or extinction memory via distinct subcortical pathways. Nature communications 13, 5540.

Yasui, Y., Breder, C.D., Saper, C.B., and Cechetto, D.F. (1991). Autonomic responses and efferent pathways from the insular cortex in the rat. The Journal of comparative neurology 303, 355–374.

